# BRN1/2 Function in Neocortical Size Determination and Microcephaly

**DOI:** 10.1101/2023.11.02.565322

**Authors:** Soraia Barão, Yijun Xu, José P. Llongueras, Rachel Vistein, Loyal Goff, Kristina Nielsen, Byoung-Il Bae, Richard S. Smith, Christopher A. Walsh, Genevieve Stein-O’Brien, Ulrich Müller

**Affiliations:** The Solomon H. Snyder Department of Neuroscience, Johns Hopkins University School of Medicine, Baltimore, MD 21205, USA; Department of Neuroscience, University of Connecticut School of Medicine, Farmington, CT 06032, USA; Northwestern University, Feinberg School of Medicine, Department of Pharmacology, Chicago, IL, 60611; Division of Genetics and Genomics, Manton Center for Orphan Disease Research, Boston Children’s Hospital, Harvard Medical School, Boston, MA 02115, USA; Howard Hughes Medical Institute, Boston Children’s Hospital, Harvard Medical School, Boston, MA 02115, USA

## Abstract

The mammalian neocortex differs vastly in size and complexity between mammalian species, yet the mechanisms that lead to an increase in brain size during evolution are not known. We show here that two transcription factors coordinate gene expression programs in progenitor cells of the neocortex to regulate their proliferative capacity and neuronal output in order to determine brain size. Comparative studies in mice, ferrets and macaques demonstrate an evolutionary conserved function for these transcription factors to regulate progenitor behaviors across the mammalian clade. Strikingly, the two transcriptional regulators control the expression of large numbers of genes linked to microcephaly suggesting that transcriptional deregulation as an important determinant of the molecular pathogenesis of microcephaly, which is consistent with the finding that genetic manipulation of the two transcription factors leads to severe microcephaly.

## Summary

The neocortex varies in size and complexity among mammals due to the tremendous variability in the number and diversity of neuronal subtypes across species^1,2^. The increased cellular diversity is paralleled by the expansion of the pool of neocortical progenitors^2–5^ and the emergence of indirect neurogenesis^6^ during brain evolution. The molecular pathways that control these biological processes and are disrupted in neurological and psychiatric disorders remain largely unknown. Here we show that the transcription factors BRN1 (POU3F3) and BRN2 (POU3F2) act as master regulators of the transcriptional programs in progenitors linked to neuronal specification and neocortex expansion. Using genetically modified lissencephalic and gyrencephalic animals, we found that BRN1/2 establish transcriptional programs in neocortical progenitors that control their proliferative capacity and the switch from direct to indirect neurogenesis. Functional studies in genetically modified mice and ferrets show that BRN1/2 act in concert with NOTCH and primary microcephaly genes to regulate progenitor behavior. Analysis of transcriptomics data from genetically modified macaques provides evidence that these molecular pathways are conserved in non-human primates. Our findings thus establish a mechanistic link between BRN1/2 and genes linked to microcephaly and demonstrate that BRN1/2 are central regulators of gene expression programs in neocortical progenitors critical to determine brain size during evolution.

An understanding of cortical progenitor diversity and of the mechanisms by which these progenitors self-renew and differentiate is critical to understand brain evolution and the defects in progenitor behavior that lead to neurological and psychiatric disorders. Cortical progenitors have been broadly divided into two classes named apical progenitors (APs) that undergo mitosis in the ventricular zone (VZ), and basal progenitors (BPs) that undergo mitosis in the subventricular zone (SVZ)^7–11^. APs engage in two modes of neurogenesis termed direct and indirect neurogenesis. During direct neurogenesis, APs divide asymmetrically to self-renew and to generate one neuron, while during indirect neurogenesis, APs divide asymmetrically to self-renew and generate a BP that then gives rise to neurons^7–11^. Indirect neurogenesis generates neurons for all cortical layers but is the predominant neurogenic mode that produces upper layer projection neurons (ULNs)^8,9,12–15^. The number and complexity of ULNs have dramatically increased in gyrencephalic brains, and indirect neurogenesis is thus intricately linked to neocortex expansion during brain evolution^3,5,16,17^. Using *in utero* electroporation and gene targeting in lissencephalic mice and gyrencephalic ferrets as well as single cell transcriptomics analysis in mice and macaques, we reveal here a central function for the POU-homeobox-domain-containing octamer-binding transcription factors BRN1 (POU3F3) and BRN2 (POU3F2) in establishing transcriptional programs that determine the proliferative capacity and neurogenic mode of cortical progenitors to regulate the production of ULNs and thus determine brain size in lissencephalic and gyrencephalic brains. Notably, BRN1 and BRN2 regulate the expression of large numbers of primary microcephaly genes thus implicating transcriptional deregulation as an important determinant of the molecular pathogenesis of microcephaly.

### BRN1/2 regulate the competence of neocortical progenitors to generate ULNs

In *Brn1/2-null* mice, the generation of ULNs is severely affected^18,19^. However, effects on brain size could not be evaluated because *Brn1/2-null* mice die at birth when the formation of cortical layers is still in progress. In addition, the molecular and cellular mechanisms by which BRN1/2 regulate neuronal specification are not well understood. Therefore, we generated mice carrying a floxed *Brn1* allele (*Brn1*^fl^) and crossed them with mice carrying a floxed *Brn2* allele (*Brn2*^fl^)^20^ to generate *Brn1^fl/fl^;Brn2^fl/fl^* mice. We then crossed *Brn1^fl/fl^;Brn2^fl/fl^*mice with *Emx1-Cre* mice^21^ on a *Brn1^fl/+^;Brn2^fl/+^* background to generate mice lacking *Brn1/2* in progenitors for excitatory neurons of the dorsal telencephalon (*Brn1/2-cKO* mice) (Extended Data Fig. 1). *Brn1/2-cKO* mice were viable. We confirmed by immunohistochemistry that inactivation of *Brn1* and *Brn2* expression followed the pattern of CRE expression and was observed in the mutant mice already at E11.5 (Extended Data Fig. 1b-e and 1h) as expected from the onset of CRE expression in *Emx1-Cre* mice around E10.5^21^. Compared to control littermates, the cortex of *Brn1/2-cKO* mice at postnatal day (P) 13 was reduced in thickness (Fig. 1a,b), had reduced cell numbers (Fig. 1c), and showed abnormal layering with cortical heterotopias (Fig. 1a). Defects in brain size were caused by aberrant generation of ULNs. Accordingly, *Brn1/2-cKO* mice lacked the corpus callosum, which is largely formed by axonal projections of ULNs (Extended Data Fig. 2a,b), and instead had an increased number of subcortical projections (Extended Data Fig. 2c,d). In the somatosensory cortex of *Brn1/2-cKO* mice, the expression of UL markers RORβ and CUX1 was absent with a concomitant increase in the numbers of cells expressing the deep layer (DL) markers TLE4 and CTIP2 (Extended Data Fig. 2e-h). The increase in DL neurons (DLNs) should have preserved at least some of the DL callosal projections but instead we observed an abnormal ventral misrouting of L1^+^ projections in *Brn1/2-cKO* mice (Extended Data Fig. 2b - open arrowhead), suggesting that axon guidance was affected in these neurons. Additionally, the number of glial cells was dramatically increased in *Brn1/2-cKO* mice at P13 (Extended Data Fig. 3). Consistent with previous studies^18,19^, *Brn1* and *Brn2* single homozygous mutants had no obvious defects in cortical development (data not shown).

**Fig. 1.**
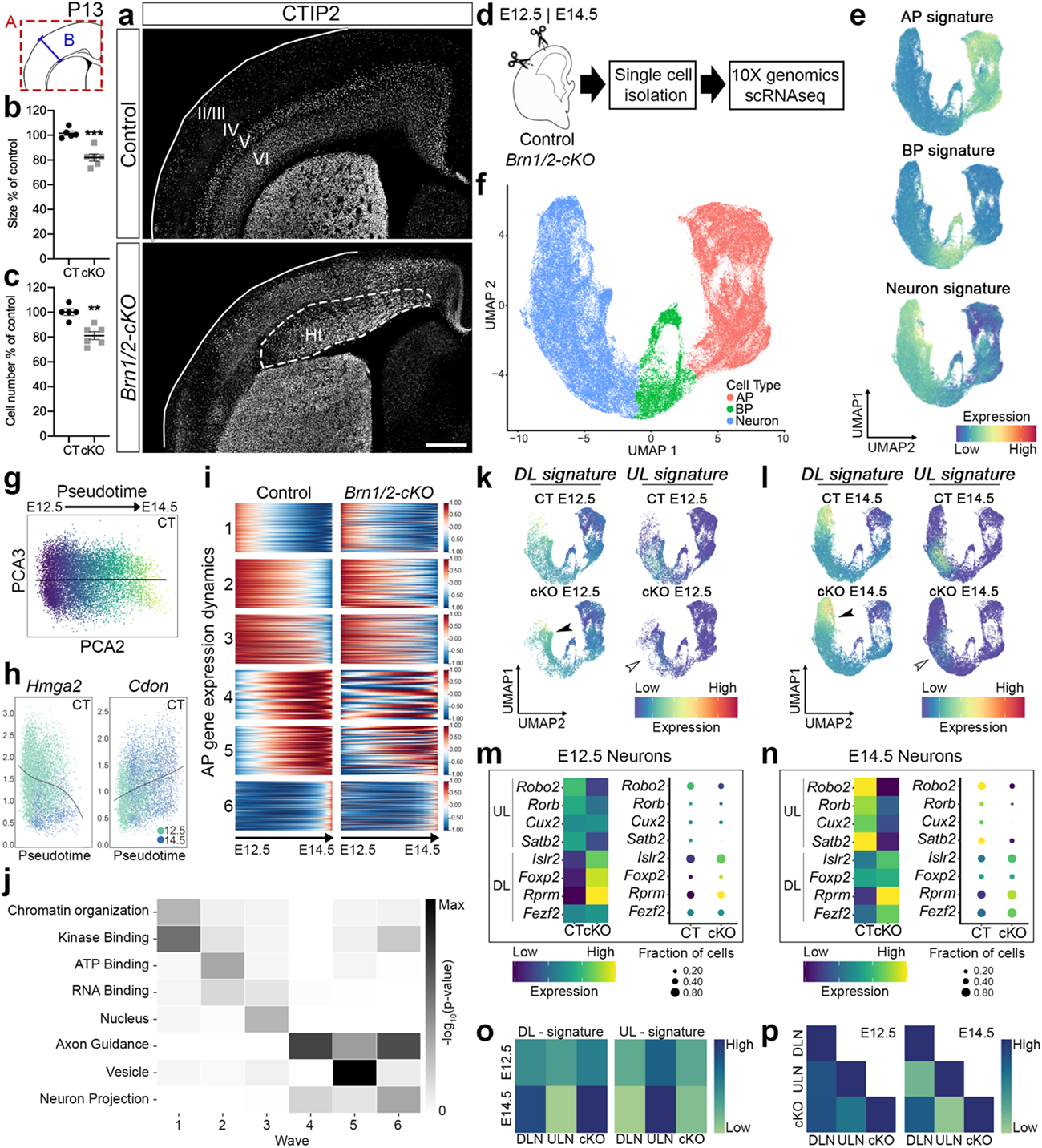
BRN1/2 regulate the competence of neocortical progenitors to generate ULNs. **a**, Control and *Brn1/2-cKO* brains analyzed by CTIP2 immunolabeling at P13. Lines represent the limits of the cortical plate (CP); dashed lines outline cortical heterotopia (Ht). **b**, **c**, Cortical size (b) and total cell number (c) in control and *Brn1/2-cKO* cortices at P13 (n=5 CT, n=6 *cKO* mice; unpaired t*-*test). **d,** 10X genomics *scRNA-seq* in control and *Brn1/2-cKO* mice at E12.5 and E14.5. **e,** UMAP of gene signatures for AP, BP and neurons. **f,** UMAPs from control and *Brn1/2-cKO* cortices at E12.5 and E14.5 by cell type. **g,** Principal component analysis (PCA) of control AP transcriptional identity organization along the pseudotime axis. **h,** Expression of *Hmga2* and *Cdon* in control APs along the pseudotime axis. **i,** Cluster analysis of the gene expression dynamics for the APs along the pseudotime axis in control and *Brn1/2-cKO* mice (1-6 represent the different transcriptomic waves along the pseudotime axis previously described by Telley et al.^27^; SI_1). **j,** Some of the most relevant gene ontology (GO) terms defining the transcriptomic waves along the pseudotime presented in i. **k, l,** UMAP of DL and UL gene signature in the scRNAseq datasets from control and *Brn1/2-cKO* cortices at E12.5 (k) and E14.5 (l). Empty arrowheads point to reduced expression of UL gene signature in mutants, arrowheads to increased expression of DL gene signature. **m, n,** Expression of the indicated cortical UL and DL marker genes in control and *Brn1/2-cKO* neurons at E12.5 (m) and E14.5 (n). **o,** Average DL and UL signature score in DLNs, ULNs and total cKO neurons at E12.5 and E14.5. **p,** Correlation of DL and UL marker gene expression among DLNs, ULNs and total cKO neurons at E12.5 and E14.5. Values are mean ± SEM; **p*<*0.01, ***p<0.001; Scale bars: 500 µm.

Prior to gliogenesis, cortical progenitors generate neurons for different cortical layers in sequential order. DLNs are generated prior to ULNs, which only begin to emerge in mice around E14.5^22^. This temporal order is largely driven by temporal changes in the competence of progenitors to generate distinct neuronal subtypes^23–29^. In addition, some progenitors for ULNs are already present at E12.5 but they begin to differentiate only after E14.5^30–33^. It has been proposed that BRN1/2 in mice are critical to regulate the transition from early-to mid-neurogenesis^18,19,34,35^, while BRN2 in monkeys already acts in early progenitors^36^. In contrast to earlier findings, we observed BRN2 expression in murine cortical progenitors already at E11.5 (Extended Data Fig. 1f,g) suggesting that BRN1/2 affect progenitor behavior in mice at earlier time points than previously thought. To test this hypothesis, we inactivated *Brn1/2* using timed *in utero* electroporation (IUE) of pCAG-CRE in *Brn1^fl/fl^;Brn2^fl/fl^* mice at E12.5 and E14.5 (Extended Data Fig. 4a). We co-expressed pCAG-RFP to identify electroporated cells. Relative to controls, BRN1/2-deficient RFP^+^ cells settled in deeper layers regardless of the time point when IUEs were performed (Extended Data Fig. 4b,c,e and f). In addition, experimental manipulations at both ages led to a reduction of BRN1/2-deficient RFP^+^ cells expressing UL markers with a concomitant increase in the number of BRN1/2-deficient RFP^+^ cells expressing DL markers (Extended Data Fig. 4b,d,e and g). Similar results were obtained when we injected EdU into pregnant mice at E12.5 and E14.5 and followed the cell-fate of the labeled progenitors by analyzing the co-labeling of EdU^+^ cells with DL and UL markers at P13 (Extended Data Fig. 4h,i,j,l and n). The EdU^+^ cells labeled at E12.5 and E14.5 also lost their preference for DLs or ULs, respectively and were distributed more broadly throughout cortical layers of *Brn1/2-cKO* mice at P13 (Extended Data Fig. 4i,j,k and m), supporting a role of BRN1/2 in neuronal migration. Altogether, these findings suggest that BRN1/2 cell-autonomously regulate the competence of progenitors to generate ULNs starting at early stages of cortical development.

To confirm this hypothesis and to determine the mechanisms by which BRN1/2 regulate neuronal specification, we performed 10X genomics single cell RNA sequencing (scRNAseq) on cortical cells isolated from *Brn1/2-cKO* and wild-type littermates at E12.5 and E14.5 (Fig. 1d). Cells were classified into cell types using cluster analysis and marker gene enrichment (Extended Data Fig. 5a). We then focused our analysis on APs, BPs and excitatory neurons (Fig. 1e,f; Extended Data Fig. 5b,c). In *Brn1/2-cKO* and controls, cell numbers were comparable (Extended Data Fig. 5d) and the proportions of AP cells decreased while the number of BPs and neurons increased between E12.5 to E14.5 (Extended Data Fig. 5d).

Telley et al. (2019) previously reported that APs undergo temporal transcriptional changes (wave 1-6) during their developmental progression^27^. We selected the most reproducible genes representing each transcriptional wave and observed in wild-type mice similar gene expression changes along the pseudotime axis as reported^27^ (Fig. 1g-j; Extended Data Fig. 5e; 1-6 represent the different transcriptomic waves along the pseudotime axis previously described by Telley et al.^27^ to reflect the temporal progression of APs and are characterized by the gene ontology (GO) terms highlighted in Fig. 1j and the full gene list available in Supplementary Information_1 (SI_1)). In contrast, the transcriptional waves were severely disrupted in APs from *Brn1/2-cKO* mice (Fig. 1i; SI_1). The transcriptional profile of E14.5 *Brn1/2-cKO* APs was more similar to the transcriptional profile of E12.5 control APs with a higher and more prolonged expression of the characteristic E12.5^high^ ^genes^ and a failure to express high levels of the characteristic E14.5^high^ ^genes^ (Fig. 1i; Extended Data Fig. 5f-j; SI_1). We also compared gene expression programs of excitatory neurons from the cortex of control and *Brn1/2-cKO* mice. The expression of UL signature genes was decreased in *Brn1/2-cKO* neurons, while the expression of DL signature genes was increased (Fig. 1k-n; Extended Data Fig. 6a,b, SI_2). Furthermore, despite the abnormal transcriptional profile of *Brn1/2-cKO* total neurons (SI_7,8), their average DL and UL signature score was closely related to the control DLNs signature score (Fig. 1o) and their gene expression strongly correlated with DL marker gene expression of control DLNs at E12.5 and E14.5 (Fig. 1p). GO terms that significantly changed in *Brn1/2-cKO* neurons included *axon guidance* and *neuron projections* (Extended Data Fig. 6c) consistent with the axonal projection defects that we observed in the mutants at later developmental time points (Extended Data Figures 2 and 6d). Pioneer neurons and glia guidepost cells were present in *Brn1/2-cKO* mice at P0 indicating the axonal projection defects observed in the mutants are primarily related to the abnormal expression of axon guidance-associated genes in these neurons at E14.5 (Extended Data Fig. 6b,d and e, SI_2). We conclude that BRN1/2 are critical for the normal regulation of temporal gene expression changes in APs that characterize their progressive restriction in fate potential to progress from the production of DLNs to ULNs.

### BRN1/2 regulate cell proliferation and the timing of cell cycle exit

To further explore the role of *Brn1/2* in progenitor function, we determined the proportion of progenitors in different stages of the cell cycle by analyzing the scRNAseq data using TRIcycle (Transferable Representation and Inference of cell cycle) (Fig. 2a). This method defines cell cycle phase based on the expression of reference genes characteristic for different stages of the cell cycle^37^: (i) G2/M, cells that are preparing to divide (G2) or already have divided (M, mitosis); (ii) S, cells that are synthesizing DNA; (iii) G1/G0, cells that are either waiting to re-enter the cell cycle (G1) or are exiting the cell cycle (G0). The number of progenitors in S-G2/M was reduced in *Brn1/2-cKO* mice and the number of progenitors in G1/G0 was increased (Fig. 2b). The differences were detected at E12.5 but were more pronounced at E14.5.

**Fig. 2.**
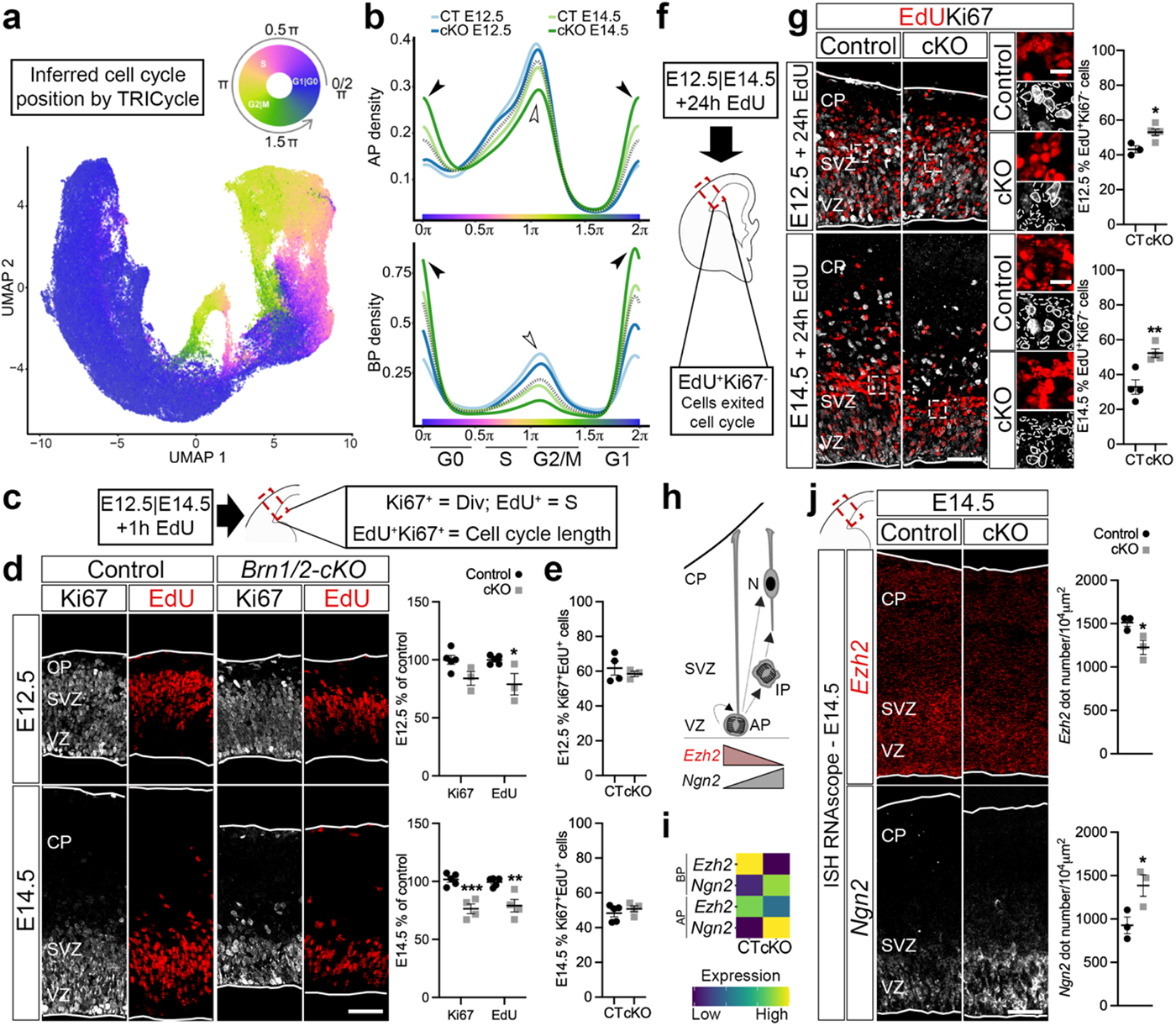
BRN1/2 regulate cell cycle exit. **a**, UMAPs from control and *Brn1/2-cKO* cortices at E12.5 and E14.5 by TRIcycle analysis^37^. **b,** Cell cycle phase analysis from control and *Brn1/2-cKO* APs and BPs at E12.5 and E14.5 represented as cell density (Wilcoxon rank sum test: AP E12.5 p= 0.085; AP E14.5 p= 1.84e-15; BP E12.5 p=1.95e-06; BP E14.5 p<2.2e-16). Empty arrowheads point to reduced S-G2/M state in mutants, arrowheads to increased G1/G0 state. **c, f,** Schematic of the experimental strategy. E12.5 and E14.5 brains were analyzed by EdU and Ki67 immunolabeling 1h (c) or 24h (f) after intraperitoneal injection of EdU. **d,** EdU (red) and Ki67 (grey) immunolabeling in control and *Brn1/2-cKO* cortical sections after 1h EdU injection at E12.5 and E14.5 (E12.5: n=5 CT, n=3 *cKO* mice; E14.5: n=5 CT, n=4 *cKO* mice; unpaired t*-*test). **e,** EdU labeling index in control and *Brn1/2-cKO* cortical sections after 1h EdU injection at E12.5 and E14.5 (E12.5: n=4 CT, n=3 *cKO* mice; E14.5: n=5 CT, n=4 *cKO* mice; unpaired t*-*test). **g,** EdU (red) and Ki67 (grey) immunolabeling in control and *Brn1/2-cKO* cortical sections after 24h EdU injection at E12.5 and E14.5 (E12.5: n=3 CT, n=5 *cKO* mice; E14.5: n=4 mice per group; unpaired t*-*test). Boxed area at higher magnification on the right. Lines and dashed lines circulating the cells show expression or absence of Ki67, respectively. **h,** Schematic of progenitors dividing and differentiating into neurons. **i,** Expression of *Ezh2* and *Ngn2* at E14.5 in control and *Brn1/2-cKO* APs and BPs as determined by scRNAseq. **j,** RNAscope for *Ezh2* (red) and *Ngn2* (grey) in control and *Brn1/2-cKO* cortical sections at E14.5 (n=3 mice per group; unpaired t*-*test). Low and top lines represent the limits of the VZ and CP, respectively. Values are mean ± SEM; \**p<*0.05, \*\**p<*0.01, ***p<0.001; Scale bars: 50 µm (lower magnification), 10 µm (higher magnification).

To analyze the extent to which progenitor behavior was altered in *Brn1/2-cKO* mice and to independently validate the TRIcycle data, we injected EdU into pregnant mice at E12.5 and E14.5 and analyzed EdU incorporation into DNA 1h later (Fig. 2c). Although the number of cortical progenitors (Extended Data Fig. 7a) and the general marker of dividing cells, Ki67 (Fig. 2d), were only significantly reduced in *Brn1/2-cKO* mice at E14.5, the number of EdU^+^ cells was reduced in *Brn1/2-cKO* mice already at E12.5 (and also at E14.5) (Fig. 2d) consistent with the reduced number of progenitors in S-G2/M we observed in mutant mice by TRIcycle analysis.

Next, since cell cycle length was unaltered in *Brn1/2-cKO* mice at E12.5 and E14.5 (Fig. 2e), we analyzed cell cycle exit by injecting EdU into pregnant mice at E12.5 and E14.5 and analyzing the number of EdU^+^/Ki67^-^ cells 24h later (Fig. 2f). The number of EdU^+^/Ki67^-^ cells was significantly increased in *Brn1/2-cKO* mice at E12.5 and E14.5 (Fig. 2g). We conclude that in *Brn1/2-cKO* mice, neocortical progenitors are proliferating less and exit the cell cycle faster. Consistent with this phenotype, the expression of numerous cell cycle-associated genes was significantly changed in the *Brn1/2-cKO* progenitors at E12.5 and at E14.5 (Extended Data Fig. 7b,c; SI_3 and SI_4). These genes included *Ezh2* and *Ngn2*, which play important roles at crucial cell cycle transition points^10,38–40^. High levels of *Ezh2* are associated with proliferation of cortical progenitors, while high levels of *Ngn2* are associated with cell cycle exit (Fig. 2h)^10,38–40^. The expression of *Ezh2* and *Ngn2* was significantly reduced and increased, respectively, in APs and BPs from *Brn1/2-cKO* mice at E14.5 (Fig. 2i; SI_3 and SI_4). We confirmed this result by ISH RNAscope for *Ezh2* and *Ngn2* (Fig. 2j). In conclusion, in the absence of BRN1/2 the proliferative capacity of progenitors is reduced, and they undergo precocious neurogenesis.

### BRN1/2 regulate the balance between direct and indirect neurogenesis via NOTCH signaling

Since ULNs are absent in *Brn1/2-cKO* mice, we wondered if BRN1/2 might determine the balance between direct and indirect neurogenesis. We combined TRIcycle with the analysis of the expression of *Tbr2* and *MKi67* (Figs. 2a and 3b). *Tbr2* is expressed in IPs and early differentiating neurons, while *MKi67* specifically labels proliferating cells. *MKi67*^+^ progenitors are thus actively dividing and among these, *Tbr*2^+^/*MKi67*^+^ defines the indirect neurogenesis branch^28^. In contrast, *MKi67*^-^ cells expressing markers for early differentiation neurons such as *Neurod2* labels primarily cells undergoing direct neurogenesis but also a small proportion of quiescent BPs and early neurons generated from IPs (Fig. 3c)^28^. The ratio between these two groups allowed us to infer the proportion of cells going through direct and indirect neurogenesis. The proportion of cells going through indirect neurogenesis was significantly decreased in *Brn1/2-cKO* mice at E12.5 and E14.5 (Fig. 3d). These findings were confirmed by TBR2 and Ki67 immunostaining (Fig. 3e) as well as by EdU incorporation analysis in TBR2^+^ cells (Extended Data Fig. 8a).

**Fig. 3.**
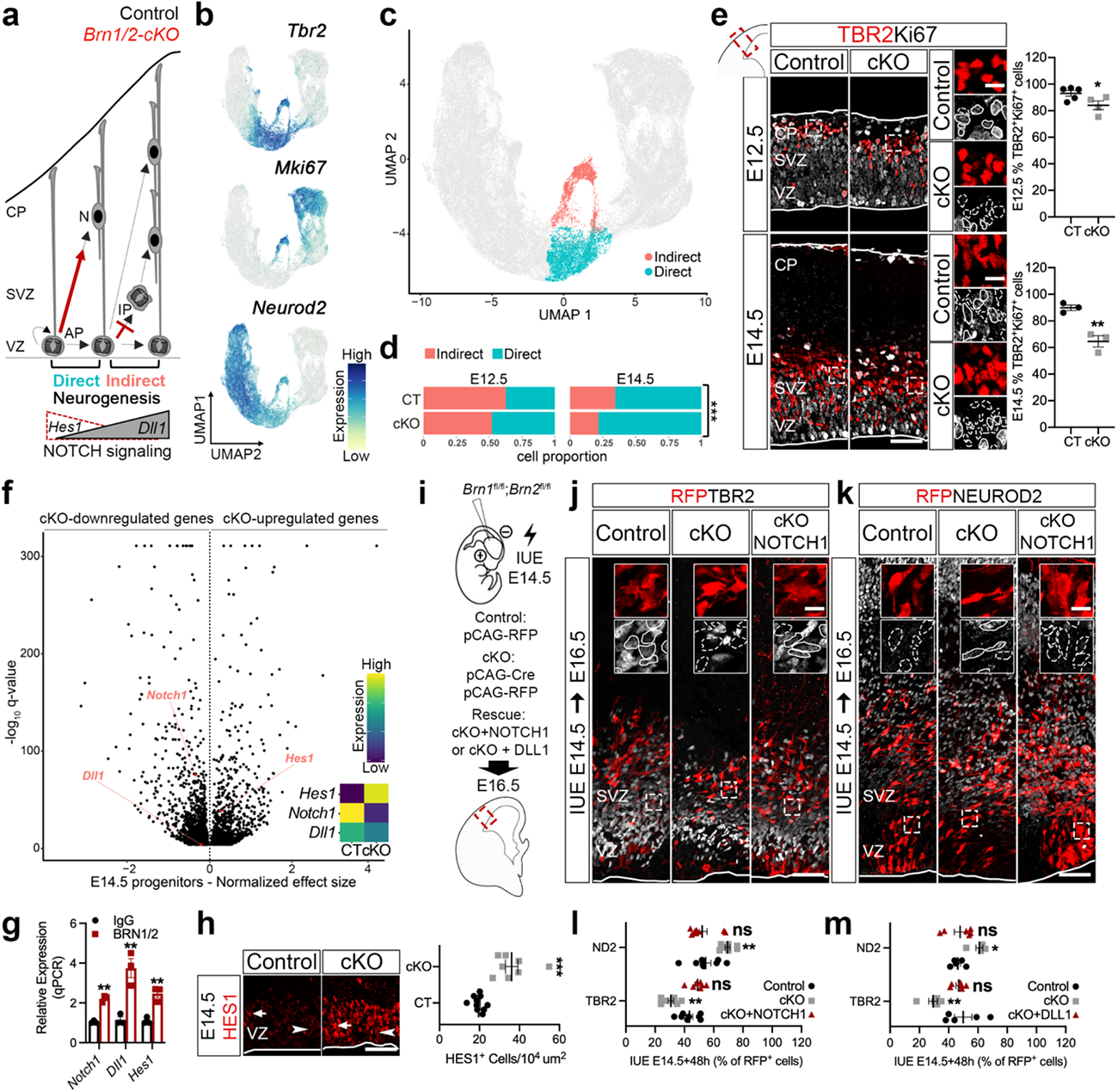
BRN1/2 regulate the switch from direct to indirect neurogenesis via NOTCH signaling. **a**, Schematic of progenitor’s neurogenic mode in control (black) and *Brn1/2-cKO* (red). **b,** UMAP of *Tbr2*, *MKi67 and Neurod2* expression in scRNAseq datasets. **c,** UMAP of cells going through indirect and direct neurogenesis in control and *Brn1/2-cKO* cortices at E12.5 and E14.5. **d,** Proportion of cells going through indirect and direct neurogenesis by age and genotype (Pearson’s Chi-squared test – ***p<0.001). **e,** TBR2 (red) and Ki67 (grey) immunolabeling in control and *Brn1/2-cKO* cortical sections at E12.5 and E14.5 (E12.5: n=5 CT, n=4 *cKO* mice; E14.5: n=3 mice per group; unpaired t*-*test). Boxed area at higher magnification on the right. Lines and dashed lines circulating cells: expression or absence of Ki67, respectively. **f,** Volcano plot: differentially expressed genes between *Brn1/2-cKO* progenitors and controls at E14.5 highlighting NOTCH signaling-associated genes (Monocle3 VGAM test; SD=0.15; q<0.05; SI_5). **g,** ChIP-qPCR analysis of BRN1/2 binding to the promoters/enhancers of the indicated genes at E14.5 (n=3 per condition; unpaired t-test). **h,** HES1 (red) immunolabeling in the VZ of control and *Brn1/2-cKO* cortical sections at E14.5 (n=10 CT, n=8 *cKO* mice; unpaired t*-*test). **i,** IUE in *Brn1^fl/fl^*;*Brn2^fl/fl^* mice at E14.5 with the indicated plasmids. **j, k,** Cell identities of the control, *Brn1/2-cKO* and *Brn1/2-cKO*+NOTCH1 condition analyzed by co-immunolabeling for RFP (red; to identify electroporated cells) with TBR2 (j, grey) and NEUROD2 (k, grey) at E16.5. Boxed area: higher magnification in inserts. Lines and dashed lines outlining cells expressing or lacking expression of the indicated marker, respectively. **l**, **m,** RFP^+^ cells expressing TBR2 and NEUROD2 at E16.5 in the control, *Brn1/2-cKO*, *Brn1/2-cKO*+NOTCH1 (l) and *Brn1/2-cKO*+DLL1 (m) condition (NOTCH1(l): n=8 CT, n=8 *cKO*, n=7 *cKO*+NOTCH1 mice; DLL1(m): n=5 CT, n=5 *cKO*, n=6 *cKO*+NOTCH1 mice; one-way ANOVA – Dunnett’s multiple comparisons test; NOTCH1-TBR2: F_2,20_=20.50; NOTCH1-NEUROD2: F_2,21_=9.702; DLL1-TBR2: F_2,13_=8.294; DLL1-NEUROD2: F_2,13_=6.748). Low and top lines represent the limits of the VZ and CP, respectively. Values are mean ± SEM; ns, not significant; *p<0.05, **p*<*0.01, ***p<0.001. Scale bars: 50 µm (lower magnification), 10 µm (higher magnification).

Similar results were obtained when we used CRE to acutely inactivate *Brn1/2* by IUE of *Brn1^fl/fl^;Brn2^fl/fl^*mice at E12.5 or E14.5 and analyzed neurogenesis at E14.5 or E16.5 (Extended Data Fig. 8b and Fig. 3i:cKO). We co-expressed pCAG-RFP to identify electroporated cells. The number of mutant RFP^+^ cells going through indirect neurogenesis was significantly reduced (Extended Data Fig. 8c and Fig. 3j,l:cKO), while the number of mutant RFP^+^/NEUROD2^+^ cells was significantly increased consistent with a higher rate of direct neurogenesis (Extended Data Fig. 8c and Fig. 3k,l:cKO). Since indirect neurogenesis is reduced already at early stages of corticogenesis, we reasoned that the generation of CUX2^+^ BPs, a subgroup of early generated BPs that is fated to generate ULNs^30,41,42^, might be affected in *Brn1/2-cKO* mice. Indeed, the proportion of *Cux2*^+^ BPs was significantly reduced in *Brn1/2-cKO* mice at E14.5 (Extended Data Fig. 8d-f), a finding that we confirmed by quantifying *Cux2* mRNA expression by ISH RNAscope in TBR2^+^ cells (Extended Data Fig. 8g). Consistent with the depletion of the CUX2^+^ BPs fated to generate ULNs in *Brn1/2-cKO* mice, scRNAseq analysis revealed that several UL markers were reduced, while some DL markers were increased in BPs from *Brn1/2-cKO* mice at E14.5 (Extended Data Fig. 8h; SI_4).

Expression levels of HES1 and the NOTCH ligand DLL1 in neural progenitors oscillate with opposite phases, which is critical to regulate the proliferation and differentiation of these cells (Extended Data Fig. 9a)^43,44^. In addition, high levels of *Hes1* and low levels of *Dll1* are associated with direct neurogenesis (Fig. 3a)^6,45^. Since, BRN1/2 suppress the activity of the *Hes1* promoter and cooperate with MASH1 to activate expression of *Dll1*^34,46^, we wondered whether defects in *Hes1* and *Dll1* expression might explain the defects in indirect neurogenesis in *Brn1/2-cKO* mice. scRNAseq analysis revealed that *Hes1* was dramatically overexpressed in *Brn1/2-cKO* progenitors at E14.5 (Fig. 3f; SI_5). In contrast, expression of *Notch1* and *Dll1* were significantly reduced (Fig. 3f; SI_5). We confirmed these findings by ISH RNAscope (Extended Data Fig. 9b). In addition, the direct binding of BRN1/2 to the *Ensembl* predicted regulatory regions of *Notch1*, *Dll1* and *Hes1* was confirmed by BRN1/2 Chromatin Immunoprecipitation at E14.5 (ChIP-qPCR; Fig. 3g), suggesting that these genes are direct targets for BRN1/2.

Given the cross-regulatory interactions between HES1 and DLL1^43,44^, we expected that the amplitude of oscillatory gene expression of NOTCH signaling pathway components might be affected in *Brn1/2-cKO* mice. We analyzed EdU incorporation and the expression of HES1 during cell cycle in control and *Brn1/2-cKO* mice 90 minutes (90’; S+G2 phase), 8h (early G1) or 14h (late G1) after EdU injection into pregnant mice at E14.5 (Extended Data Fig. 9c). Although the typical HES1 oscillations still occurred in *Brn1/2-cKO* progenitors (Extended Data Fig. 9c), levels of HES1 were vastly increased compared to controls at any time during the cell cycle (Fig. 3h; Extended Data Fig. 9c). Even lowest levels of HES1 in mutants never fell below highest levels in controls (Fig. 3h; Extended Data Fig. 9c). We conclude that BRN1/2 play a crucial role in establishing normal expression patterns of NOTCH signaling pathway components in progenitors.

To provide direct evidence for a causal relationship between perturbations in NOTCH signaling and the changes in indirect neurogenesis in *Brn1/2-cKO* mice, we co-expressed pCAG-CRE to inactivate *Brn1/*2 and pCS2-NOTCH1 to overexpress NOTCH1 by IUE of *Brn1^fl/fl^;Brn2^fl/fl^*mice at E14.5 and analyzed neurogenesis at E16.5 (Fig. 3i). We co-expressed pCAG-RFP to identify electroporated cells. While NOTCH1 overexpression did not change the rate of indirect neurogenesis (RFP^+^/TBR2^+^ cells) and neuronal production (RFP^+^/NEUROD2^+^ cells) in wild-type (Extended Data Fig. 9d-f), it restored levels of indirect neurogenesis and neuronal production in the *Brn1/2-cKO* condition to control levels (Fig. 3j-l). Significantly, overexpression of DLL1 in *Brn1^fl/fl^;Brn2^fl/fl^* mice phenocopied the effect of NOTCH1 overexpression (Fig. 3m; Extended Data Fig. 9g-i), which is consistent with the idea that an imbalance in DLL1-dependent NOTCH1 signaling in progenitors is responsible for the defects in indirect neurogenesis in *Brn1/2-cKO* mice (Fig. 3a).

### BRN1/2 are required to maintain the neuronal progenitor pool through the regulation of microcephaly-associated genes

Increases in brain size during evolution are accompanied by a disproportionate increase in ULNs compared to DLNs, and the production of ULNs is frequently affected in microcephaly patients^17^. Because ULNs are massively affected by defects in BRN1/2 function and the brain of *Brn1/2-cKO* mice is vastly more microcephalic than is typical for most microcephaly mouse models, we wondered whether BRN1/2 are required for the expression and function of sets of genes linked to microcephaly. ScRNAseq analysis revealed that the expression of a large number of genes that when mutated cause microcephaly in humans was significantly decreased in *Brn1/2-cKO* progenitors (Fig. 4a; SI_5). These included 11 classical primary microcephaly genes: *Cdk5rap2* (MCPH3), *Knl1* (MCPH4), *Aspm* (MCPH5), *Cenpj* (MCPH6), *Stil* (MCPH7), *Cep135* (MCPH8), *Cdk6* (MCPH12), *Sass6* (MCPH14), *Mfsd2a* (MCPH15), *Cit* (MCPH17) and *Copb2* (MCPH19)^47^. We confirmed these findings for *Aspm* and *Cdk6* by ISH RNAscope at E14.5 (Extended Data Fig. 10a).

**Fig. 4.**
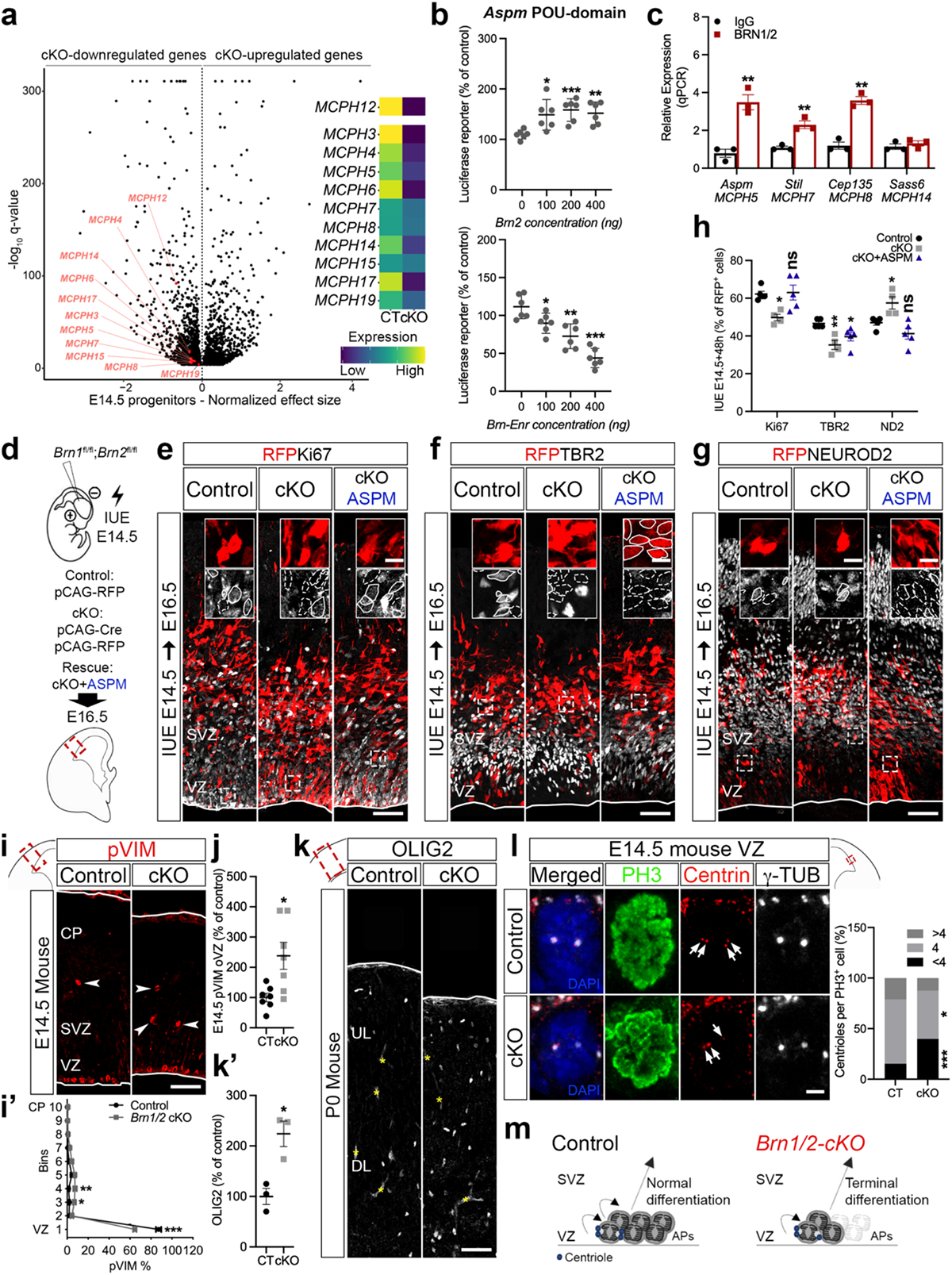
BRN1/2 are required for the expression of microcephaly-associated genes and maintenance of the neuronal progenitor pool. **a**, Volcano plot: genes differentially expressed between *Brn1/2-cKO* and control progenitors at E14.5 highlighting primary microcephaly genes (Monocle3 VGAM test; SD=0.15; q<0.05; SI_5). **b,** Luciferase reporter-activity of *Aspm* gene regulator region when co-expressed with different concentrations of *Brn2* or *Brn-Enr* in HEK 293 cells (n=6 per tested condition; unpaired t-test). **c,** ChIP-qPCR analysis of BRN1/2 binding to the promoters/enhancers of the indicated genes at E14.5 (n=3 per condition; unpaired t-test). **d,** IUE in *Brn1^fl/fl^*;*Brn2^fl/fl^* mice at E14.5 with the indicated plasmids. **e, f, g,** Cell identities of the control, *Brn1/2-cKO* and *Brn1/2-cKO*+ASPM condition analyzed by co-immunolabeling for RFP (red; to identify electroporated cells) with Ki67 (e, grey), TBR2 (f, grey) and NEUROD2 (g, grey) at E16.5. Boxed area: higher magnification in inserts. Lines and dashed lines outlining cells expressing or lacking expression of the indicated marker, respectively. **h,** RFP^+^ cells expressing TBR2 and NEUROD2 at E16.5 in the control, *Brn1/2-cKO* and *Brn1/2-cKO*+ASPM condition (n=5 CT, n=4 *cKO*, n=5 *cKO*+ASPM mice; one-way ANOVA – Dunnett’s multiple comparisons test; Ki67: F_2,11_=6.582; TBR2: F_2,11_=8.681; NEUROD2: F_2,11_=10.02). **i,** pVIM immunolabeling in cortical sections of control and *Brn1/2-cKO* mice at E14.5. Arrowheads: pVIM^+^ cells outside the VZ. **i’**, Distribution of pVIM^+^ cells in the cortex of control and *Brn1/2-cKO* mice (n=7 mice per group; two-way ANOVA – Šídák’s multiple comparisons test; F_9,120_=25.73). **j,** pVIM^+^ cells outside of the VZ (oVZ) in control and *Brn1/2-cKO* cortices (n=7 mice per group; unpaired t-test). **k, k’,** OLIG2 immunolabeling in cortical sections of control and *Brn1/2-cKO* mice at P0 (n=3 mice per group; unpaired t-test). **l,** Centriole number per PH3^+^ cell in the VZ of control and *Brn1/2-cKO* mice at E14.5 (n=3 mice per group; unpaired t-test). Arrows indicate the centrioles. **m,** Schematic of centrosome function and progenitor cell differentiation in control (black) and *Brn1/2-cKO* (red). Low and top lines represent the limits of the VZ and CP, respectively. Yellow asterisks indicate auto-fluorescent blood vessels. Values are mean ± SEM; \**p<*0.05, \*\**p<*0.01, ***p<0.001. Scale bars: 50 µm (lower magnification; e-g, i, and k), 10 µm (higher magnification; e-g), 2 µm (l).

To study the regulation of *Aspm* by BRN1/2, we cloned the *Ensembl* predicted regulatory region of *Aspm* containing POU-domain binding sites in a luciferase reporter vector and tested its transcriptional activity in absence or presence of pCAG-BRN2 or pCAG-*Brn*-DBD-EnR, a dominant negative inhibitor of BRN1/2 function^34,46^. Co-expression with BRN2 increased *Aspm*-luciferase activity while co-expression with *Brn*-DBD-EnR inhibited *Aspm*-luciferase activity (Fig. 4b) indicating BRN1/2 directly regulates the expression of *Aspm*. The direct binding of BRN1/2 to the *Ensembl* predicted regulatory regions of *Aspm* was confirmed by ChIP-qPCR at E14.5 (Fig.4c; Extended Data Fig. 10b). To understand the extent to which *Aspm* contributes to the changes of progenitor behavior in *Brn1/2-cKO* mice, we co-expressed pCAG-CRE to inactivate *Brn1/*2 and pBlue-hASPM to overexpress ASPM by IUE of *Brn1^fl/fl^;Brn2^fl/fl^*mice at E14.5 and analyzed neurogenesis at E16.5 (Fig. 4d). We co-expressed pCAG-RFP to identify electroporated cells. While ASPM overexpression did not rescue the rate of indirect neurogenesis (RFP^+^/TBR2^+^ cells; Fig. 4f,h), it restored the levels of proliferation/cell cycle exit (RFP^+^/Ki67^+^ cells) and neuronal production (RFP^+^/NEUROD2^+^ cells) in the *Brn1/2-cKO* condition to control levels (Fig. 4e, g and h). These results indicate that BRN1/2-dependent regulation of *Aspm* expression and function primarily controls the proliferation and cell cycle exit of progenitor cells without major contribution to the levels of indirect neurogenesis that are mainly regulated by the levels of NOTCH signaling in these cells.

Knock-out of *ASPM* in ferret, a small carnivore characterized by an expanded and gyrencephalic neocortex, phenocopies the microcephaly defect in human patients^48^. To gain insights on the cellular mechanisms controlled by BRN1/2-dependent regulation of *Aspm* expression, we compared the cortical phenotype of *Brn1/2-cKO* mice and *ASPM*KO ferrets. In *ASPM*KO ferrets, APs detached prematurely from the VZ resulting in increased number of terminal divisions and premature differentiation towards the glial lineage (Extended Data Fig. 10c-e and h-j)^48^. Although the distribution of TBR2^+^Ki67^+^ cells within the neocortex was broadened including localization of cells outside of the VZ (Extended Data Fig. 10f), the levels of indirect neurogenesis were not affected in *ASPM*KO ferrets at E35 (Extended Data Fig. 10g), consistent with the IUE results where ASPM overexpression failed to rescue the balance between direct and indirect neurogenesis in *Brn1/2-cKO* mice. Similar to *ASPM*KO ferrets, APs in *Brn1/2-cKO* mice were displaced at E14.5 (Fig. 4i,j) and the number of glial cells was also increased at P0 (Fig. 4k). In addition, the number of neuronal progenitors declined more rapidly throughout embryonic development in *Brn1/2-cKO* mice compared to controls (Extended Data Fig. 10k) and glial numbers were further increased at P13 (Extended Data Fig. 3a).

To further determine if there is a precocious shift from neurogenesis to gliogenesis in *Brn1/2-cKO* mice comparable to *ASPM*KO, we injected EdU into pregnant mice at E12.5 and E14.5 and followed the cell-fate of the labeled progenitors by analyzing the co-labeling of EdU^+^ cells with the glia markers SOX9 and OLIG2 at P13 (Extended Data Fig. 3b). Numbers of EdU^+^SOX9^+^ and EdU^+^OLIG2^+^ cells were significantly increased in *Brn1/2-cKO* mice at P13 (Extended Data Fig. 3c-f), which is indicative of precocious gliogenesis happening in *Brn1/2-cKO* mice during early embryonic development. We obtained similar results when we inactivated *Brn1/2* in the developing neocortex of *Brn1^fl/+^;Brn2^fl/+^* by IUE of pCAG-CRE at E14.5 followed by the quantification of the numbers of GFP^+^OLIG2^+^ cells at P13 (Extended Data Fig 3g-i).

Since *Aspm* and most of the microcephaly-associated genes altered in *Brn1/2-cKO* progenitors encode centrosome proteins^47^, we tested how the absence of BRN1/2 affects centrosome function in progenitors. The expression of genes essential for centrosome function were significantly reduced in *Brn1/2-cKO* progenitors at E14.5 (Extended Data Fig. 10l; SI_5) and centriole duplication was disrupted (Fig. 4l) indicating that defects in *ASPM* expression, and possibly in other microcephaly genes like *Stil* and *Cep135*, contribute to the cortical phenotype of *Brn1/2-cKO* mice (Fig. 4m).

### BRN1/2 function is conserved across phylogeny

In *BRN2*KO monkeys, the neurogenic behavior of cortical progenitors is similarly altered as in *Brn1/2-cKO* mice^36^. To test if the mechanisms by which BRN1/2 regulate progenitor behavior are evolutionary conserved, we first inhibited BRN1/2 activity in wild-type ferrets at E35 by IUE with pCAG-*Brn*-DBD-EnR (Fig. 5a)^34,46^. 48h after *Brn*-DBD-EnR expression, the number of dividing progenitors (RFP^+^/Ki67^+^) was significantly reduced, indicative of faster cell cycle exit compared to controls (Fig. 5b,e). The number of cells going through indirect neurogenesis (RFP^+^/TBR2^+^) was significantly reduced (Fig. 5c,e), while neuronal output (RFP^+^/NEUROD2^+^ cells) was increased (Fig. 5d,e). Additionally, the expression of *Notch1* was reduced in the cortex of ferrets expressing *Brn*-DBD-EnR (Fig. 5f-f’). Thus, our results in ferrets mirror those in mice, providing evidence for a conserved role for BRN1/2 in cortical progenitors of lissencephalic and gyrencephalic brains. In further support of this hypothesis, in the cortex of ferrets expressing *Brn*-DBD-EnR the expression of *Aspm* and *Cdk6* was also reduced (Fig. 5g), resulting in an increase of RFP^+^/OLIG2^+^ cells and total OLIG2^+^ cells (Fig. 5h,h’), indicating a precocious shift from neurogenesis to gliogenesis in ferrets in absence of BRN1/2 activity.

**Fig. 5.**
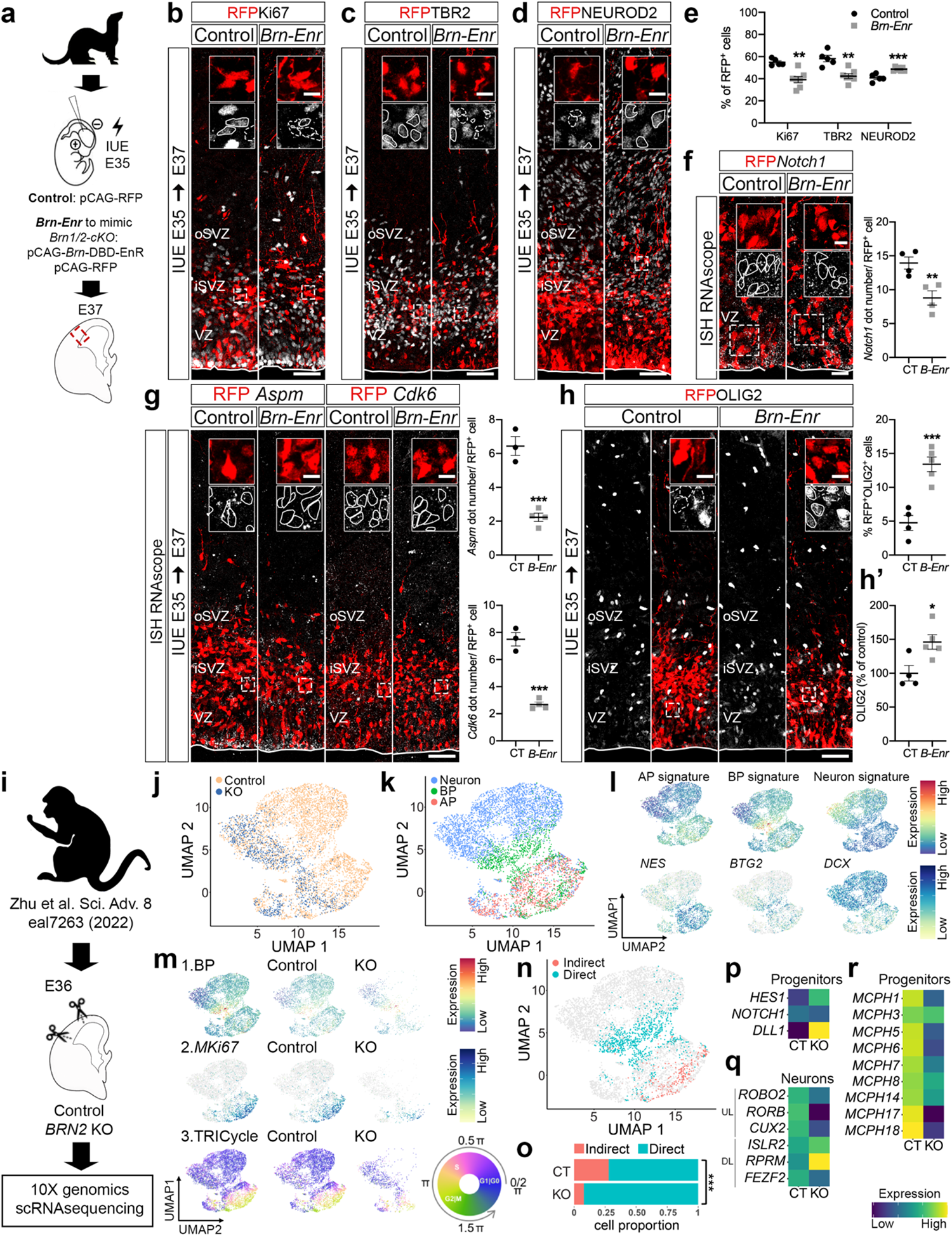
BRN1/2 function is conserved across mammalian species. **a**, IUE in wild-type ferrets at E35 with the indicated plasmids. **b-d**, Cell identities in the control and *Brn-Enr* condition analyzed by co-immunolabeling of RFP (red) with Ki67 (b, grey), TBR2 (c, grey) and NEUROD2 (d, grey) at E37. Boxed area: higher magnification in inserts. Lines and dashed lines outline cells expressing or not expressing the indicated marker, respectively. **e,** RFP^+^ cells expressing Ki67, TBR2 and NEUROD2 in the electroporated somatosensory cortex (n=4 CT, n=7 *Brn-Enr* ferrets; unpaired t*-* test). **f,** *Notch1* (grey) expression by RNAscope and RFP (red) expression by immunolabeling in the VZ of the electroporated somatosensory cortex. Boxed area: higher magnification in inserts. Lines outline RFP^+^ cells quantified for *Notch1* mRNA expression (n=4 ferrets per group; unpaired t*-*test). **g,** *Aspm* or *Cdk6* (grey) expression by RNAscope and RFP (red) expression by immunolabeling in the electroporated somatosensory cortex. Boxed area: higher magnification in inserts. Lines outline RFP^+^ cells quantified for *Aspm* or *Cdk6* mRNA expression (n=3 CT, n=4 *Brn-Enr* ferrets; unpaired t*-*test). **h,** RFP^+^ cells expressing OLIG2 (gray) in the electroporated somatosensory cortex (n=4 CT, n=5 *Brn-Enr* ferrets; unpaired t*-*test). Boxed area: higher magnification in inserts. Lines and dashed lines circulating cells: expression or absence of OLIG2, respectively. **h’,** Total OLIG2^+^ cells in the electroporated somatosensory cortex (n=4 CT, n=5 *Brn-Enr* ferrets; unpaired t*-*test). **i,** ScRNAseq DATA from E36 cortex of control and *BRN2*KO cynomolgus monkey^36^ re-analyzed using the same pipeline we used for the analysis in mice. **j**, **k,** UMAP of scRNAseq from control and *BRN2*KO cortices by genotype (j) and cell type (k). **l,** UMAP of gene signatures for AP (e.g., *NES*), BP (e.g., *BTG2*) and neurons (e.g., *DCX*) in control and *BRN2*KO cortices. **m,** UMAP of BP signature (1), *MKi67* expression (2) and TRIcycle score (3) in control and *BRN2*KO cortices. **n,** UMAP of cells going through indirect and direct neurogenesis in control and *BRN2*KO cortices. **o,** Proportion of cells going through indirect and direct neurogenesis by genotype (Pearson’s Chi-squared test – ***p<0.001). **p,** Expression of NOTCH signaling-associated genes in control and *BRN2*KO progenitors. **q,** Expression of the indicated cortical UL and DL neuronal marker genes in control and *BRN2*KO neurons. **r,** Expression of primary microcephaly genes differentially affected in *BRN2*KO progenitors compared to controls. Values are mean ± SEM; *p<0.05, **p<0.01, ***p<0.001. Scale bars: 50 µm (lower magnification), 10 µm (higher magnification).

Next, we analyzed available raw scRNAseq data from *BRN2*KO monkeys at E36^36^ using identical criteria as employed in our analysis of scRNAseq from mice (Fig. 5i-k). We identified APs, BPs and neurons (Fig. 5l) and used the BP gene expression signature, the expression profile of *MKi67* and TRIcycle analysis (Fig. 5m1-3), to identify cells going through direct and indirect neurogenesis (Fig. 4n). Similar to our findings in mice, the proportion of cells going through indirect neurogenesis was significantly reduced in *BRN2*KO monkeys (Fig. 5o). Expression of *NOTCH1* was reduced while *HES1* was dramatically increased in *BRN2*KO progenitors (Fig. 5p; SI_6). Unlike in mice, *DLL1* expression was increased in *BRN2*KO progenitors (Fig. 5p; SI_6). Effects on the expression of *DLL1* might not be observed in monkeys because BRN1 may compensate in part for loss of BRN2. Alternatively, the regulation of NOTCH signaling pathways in monkeys might reflect differences between species. Interestingly, the expression of some UL markers was already reduced in the *BRN2*KO neurons at E36 (Fig. 5q; SI_6). In addition, the expression of 9 classical microcephaly genes (*MCPH1, CDK5RAP2* (MCPH3), *ASPM* (MCPH5), *CENPJ* (MCPH6), *STIL* (MCPH7), *CEP135* (MCPH8), *SASS6* (MCPH14), *CIT* (MCPH17) and *WDFY3* (MCPH18))^47^ was reduced in *BRN2*KO progenitors at E36 (Fig. 5r; SI_6) mimicking the results in *Brn1/2-cKO* mice.

Overall, our findings reveal a striking similarity in the defects caused by perturbations in BRN1/2 function in mice, ferrets and monkeys, supporting a model in which BRN1/2 have evolutionarily conserved functions to regulate neocortical progenitor behavior in lissencephalic and gyrencephalic animals.

## Discussion

The molecular mechanisms that regulate the behavior of cortical progenitors in the mammalian clade and thus control brain size are not well understood. We show here that BRN1/2 are key drivers of the transcriptional programs in neocortical progenitors of mice, ferrets and monkeys that determine the timing of their differentiation and their total output of ULNs. Loss of BRN1/2 in progenitors leads to the deregulation of the expression of genes that regulate cell cycle progression, NOTCH signaling and centrosome function, thus coordinating the expression of sets of genes that affect different aspects of progenitor behavior.

We show that BRN1/2 are expressed in cortical progenitors of mice already at E11.5 during lower layer neurogenesis. BRN2 in monkeys is also expressed in cortical progenitors from the earliest stages of neurogenesis^36^, suggesting evolutionary conservation in the regulation of BRN2 expression. Consistent with the BRN1/2 expression pattern, we demonstrate that BRN1/2 act in progenitors during the earliest stages of cortical development to regulate their transcriptional programs, thus leading to defects in cell proliferation, precocious cell cycle exit, and altered balance of direct and indirect neurogenesis.

Our functional studies demonstrate that deregulation of NOTCH signaling contributes to defects in indirect neurogenesis and the production of ULNs observed in *Brn1/2-cKO* mice, which is consistent with previous studies demonstrating the crucial function of NOTCH signaling for regulating the neurogenic mode of cortical progenitors^6,45^. We show that BRN1/2 directly bind to the predicted regulatory regions of *Notch1*, *Dll1* and *Hes1* promoters/enhancers. Consistent with this finding, we observed increased levels of *Hes1* and reduced levels of *Dll1* and *Notch1* in *Brn1/2-cKO* mice. Expression of NOTCH signaling components in wild-type mice is tightly regulated by feedback mechanisms that establish their oscillatory expression and the amplitude of these oscillations is crucial for balanced levels of progenitor’s proliferation and differentiation^43,44^. Oscillations at higher levels than normal have been shown to inhibit proliferation^49,50^. In *Brn1/2-cKO* mice, the lowest HES1 expression levels during cell cycle are higher than peak levels of HES1 in wild-type, consistent with the proliferative defects in the mutant mice. Also, *Dll1* and *Hes1* are intricately linked to the neurogenic mode of cortical progenitors, where low levels of *Dll1* are associated with reduced indirect neurogenesis^6^, while high levels of *Hes1* are associated with high levels of direct neurogenesis^45^, which is consistent with the phenotype of *Brn1/2-cKO* mice. Remarkably, BRN1/2 functional defects are restored when we overexpress NOTCH1 or DLL1 to override the loss of BRN1/2 function, indicating that NOTCH signaling components regulate the neurogenic mode of cortical progenitors downstream of BRN1/2.

In addition to the transcriptional changes in cell cycle-and NOTCH signaling-associated genes, the expression of large numbers of microcephaly-associated genes is altered in *Brn1/2-cKO* mice. In addition, we show that BRN1/2 directly bind and regulate the transcriptional activity of *Aspm*, a gene essential for normal centrosome function. Centrosomes have crucial roles in the organization of microtubules and regulate cell polarity, the formation of primary cilia and mitotic spindle assembly^51^. We found that centrosome function is disrupted in *Brn1/2-cKO* progenitors. Consistent with the deregulation of the expression of microcephaly genes as shown by perturbation of ASPM function^48^, defects in centrosome function result in precocious cell cycle exit in *Brn1/2-cKO* progenitors and accelerated depletion of neuronal progenitors during progressive stages of neurogenesis, contributing to the absence of ULNs, increased gliogenesis and reduced brain size.

Interestingly, the *Brn1/2-cKO* proliferation defects but not the levels of indirect neurogenesis are restored when we overexpress ASPM to override the loss of BRN1/2 function, indicating that BRN1/2-dependent regulation of *Aspm* expression and function primarily controls the proliferation and cell cycle exit of progenitor cells without major contribution to the levels of indirect neurogenesis that are mainly controlled by the tightly regulated NOTCH signaling by BRN1/2 in these cells.

While our findings provide evidence for an evolutionary conserved role for BRN1/2 regulation of transcriptional programs that control progenitor behavior, it remains to be established to what extent the function of these transcriptional regulators has been modified during brain evolution to produce differences in brain size. For example, disruption of BRN2 function alone affects cortical development in monkeys^36^, but not in mice^18,19^, suggesting that redundancy of BRN1 and BRN2 function is not universally conserved. In addition, recent studies have shown that during evolution, enhancers that control neuronal gene expression have been altered by mutations that affect predominantly binding sites for a limited set of transcription factors, including BRN1/2^52^. Microcephaly-associated genes are included in the list of genes that contain enhancer elements enriched in BRN1/2 binding sites^52^. Changes to regulatory regions of the genome that affect the transcriptional landscape are thus likely to be intricately linked to adaptive changes crucial for neocortical expansion and affected in microcephaly in humans.

## Methods

### Mice

All animal experiments adhered to the guidelines of the National Institute of Health and were approved by the Institutional Animal Care and Use Committee at Johns Hopkins University School of Medicine. Mice were maintained on a 14h light/10h dark cycle. Both male and female mice were used, and no obvious differences between the sexes were noted. All mice were group-housed in pathogen-free facilities with regulated temperature and humidity and given *ad libitum* access to food and water. All mice used were seemingly free of infection, health abnormalities or immune system deficiencies and were employed independently of their gender. None of the mice used had been used for previous experiments. The date of the vaginal plug detection was designated E0.5, and the date of birth P0. *Emx1-Cre* [B6.129S2-*Emx1tm1(cre)Krj*]^21^ and *Brn2* conditional knockout [*Brn-2^fl/fl^*]^20^ mice have been described. *Brn1* conditional knockout mouse was generated in collaboration with Janelia Research Campus - Howard Hughes Medical Institute. The embryonic stem cells (ESCs) were designed to insert loxP sites flanking the coding sequence of *Brn1* gene, with upstream elements including a neomycin-resistance cassette (PGK-neo) and *lacZ* reporter flanked by two FRT sites. ESCs were transplanted into mouse blastocysts to produce transgenic mice. Heterozygous F1 mice (*Brn1flox-neo/+*) were mated with *FLPe* mice to remove the PGK-neo cassette and *lacZ* reporter. The resulting offspring were subsequently mated to C57BL/6J mice to remove the *FLPe* transgene. C57BL/6J wild type mice were obtained from Charles River Laboratories. *Brn1^fl/fl^* and *Brn2^fl/fl^*mice were crossed and the resulting *Brn1^fl/fl^;Brn2^fl/fl^*mice were crossed with heterozygous *Emx1-Cre* mice. Animals with no CRE-recombination were used as control littermates in the scRNAseq experiments. Animals with no CRE-recombination or heterozygous for one or the two genes (*Brn1* and *Brn2*) were used as control littermates in the different histological experiments. We used these mice interchangeably because they were indistinguishable from each other and from wild-type mice.

### Ferrets

All procedures adhered to the guidelines of the National Institute of Health and were approved by the Animal Care and Use Committee at Johns Hopkins University. Experiments were performed in female ferrets (*Mustela putoris furo,* Marshall BioResources) with normal immune status. Animals were housed in a 16h light/8h dark cycle. Ferrets were not involved in previous studies. The *ASPM*KO brain sections^48^ were obtained through the collaboration with Dr. Byoung-Il Bae, Dr. Richard S. Smith and Dr. Christopher A. Walsh.

### Immunohistochemistry

Embryonic brains were fixed in 4% paraformaldehyde (PFA) in phosphate-buffered saline (PBS) for 2-4h at 4°C. Postnatal mice were transcardially perfused with 20 ml ice-cold 4% PFA using a peristaltic pump at a rate of 2 ml/min. Brains were removed from the skull and postfixed in 4% PFA overnight at 4°C. Embryonic brains were cryopreserved with 30% sucrose, embedded in Tissue-Tek O.C.T. compound (#4583, SAKURA) and frozen at -80°C. They were then sectioned coronally at 16 µm with a cryostat (CM3050 S, Leica). Postnatal brains were embedded in 3% low melting point agarose (#R0801, Thermo Fisher Scientific) in PBS and sectioned coronally or sagittally at 60 µm with a vibrating microtome (VT1200S, Leica). For immunohistochemistry, brain sections were permeabilized in PBS containing 0.2% Triton X-100 and blocked in 10% goat serum for 1h at room temperature. Brain slices were then incubated with primary antibodies in blocking solution overnight (or 48h depending on the thickness of the section and antibody used) at 4°C and subsequently incubated with the appropriate secondary antibodies diluted in blocking solution for 2h at room temperature. To label all cell nuclei, the fluorescent nuclear dye DAPI (1 μg/ml, #D9542, Sigma-Aldrich) was included with the secondary antibody solution. The sections were mounted with ProLong^TM^ Gold (#P36930, Thermo Fisher Scientific). Primary antibodies used were: anti-L1 (1:500, rat monoclonal, MAB5272 Millipore Sigma); anti-NRP1 (1:500, goat polyclonal, AF566 R&D systems); anti-TLE4 (1:500, rabbit polyclonal, ab64833 Abcam); anti-CTIP2 (1:1000, rat monoclonal, ab18465 Abcam); anti-RORβ (1:500, mouse monoclonal, PP-N7927-00 R&D systems); anti-CUX1 (1:1000, rabbit polyclonal, 11733-1-AP Proteintech); anti-GFP (1:500, chicken polyclonal, GFP-1020 Aves Labs Inc.); anti-BRN2 (1:1000, rabbit polyclonal, PA5-30124 Thermo Fisher Scientific or 1:1000, rabbit monoclonal, 12137 Cell Signaling); anti-BRN1 (1:1000, rabbit polyclonal, sc-6028-R Santa Cruz Biotechnology); anti-CRE (1:200, mouse monoclonal, MAB3120 Millipore Sigma); anti-NeuN (1:500, mouse monoclonal, MAB377 Millipore Sigma); anti-Ki67 (1:500, rabbit polyclonal, ab15580 Abcam or 1:500, rat monoclonal, 14-5698-82 Thermo Fisher Scientific); anti-RFP (1:1000, rat monoclonal, 5F8 ChromoTek or 1:1000, rabbit polyclonal, ab62341 Abcam); anti-Pax6 (1:250, mouse monoclonal, MA1-109 Thermo Fisher Scientific); anti-TBR2 (1:500, rat monoclonal, 14-4875-82 Thermo Fisher Scientific or 1:500, rabbit polyclonal, ab183991 Abcam); anti-NEUROD2 (1:500, rabbit polyclonal, ab104430 Abcam); anti-HES1 (1:1000, rabbit polyclonal, R. Kageyama gift^43^); anti-pVIM (1:500, mouse monoclonal, D076-3 MBL); anti-OLIG2 (1:500, rabbit polyclonal, AB9610 Millipore Sigma); anti-SOX9 (1:1000, rabbit monoclonal, ab185966 Abcam); anti-GFAP (1:1000, chicken polyclonal, AB5541 Millipore Sigma); anti-PH3 (1:1000, rat monoclonal, ab10543 Abcam); anti-γ-TUBULIN (1:500, goat polyclonal, A. Holland gift^53^); anti-Centrin (1:500, rabbit polyclonal, A. Holland gift^54^). Secondary antibodies used were: Alexa 555, 647, 488 anti-rat (A21434; A21247, A11006), anti-rabbit (A21430; A21246) and anti-mouse (A21425; A21237); Alexa 488 anti-rabbit (A11070); Alexa 488 and 647 anti-chicken (A11039; A21449); anti-rabbit HRP (A27036); all diluted 1:500 and all from Thermo Fisher Scientific. Antigen retrieval^55^ was performed for all immunohistochemistry at embryonic ages and to stain for OLIG2 and RORβ postnatally. The method used here was a heat-induced citrate method^55^. HES1 immunostaining was done using a horseradish peroxidase (HRP)-secondary antibody and an additional tyramide signal amplification (TSA) step. Therefore, after antigen retrieval, the brains sections were treated with 3% hydrogen peroxide to block HRP-unspecific binding. After the secondary antibody, the sections were washed and incubated with TSA solution (#NEL744001KT, Akoya Biosciences) overnight at 4°C. The sections were then mounted with ProLong^TM^ Gold.

### RNAscope multiplex *in situ* hybridization

Brain sections were processed for multiplex fluorescent *in situ* hybridization RNAscope^56^ following the manufacturer’s instructions (#323110, Advanced Cell Diagnostics). Gene-specific probes: *Fezf2* (313301-C2), *Satb2* (413261), *Ezh2* (446611-C2), *Ngn2* (417291-C2), *Cux2* (469551-C3), *Notch1* (404641), *Dll1* (425071-C2)*, Hes1* (417701)*, Aspm* (427881) *and Cdk6* (570091-C2). When we combined RNAscope with immunohistochemistry, the sections were briefly washed in 1X PBS after the last conjugation step and the immunohistochemistry protocol was followed as previously described.

### EdU injections and labeling

EdU (5-ethynyl-2’-deoxyuridine; #A10044, Thermo Fisher Scientific) was diluted at 10mg/ml in 0.9% NaCl (#114-055-101, Quality Biological) and administered in pregnant females at the specified embryonic pregnancy stage at 50mg/Kg body weight. For proliferation and cell cycle length analysis, a single intraperitoneal EdU injection was administered at E12.5 or E14.5, embryos were fixed after 1h and the number of EdU^+^ cells and the percentage of Ki67^+^ cells labeled with EdU were quantified. For cell cycle exit calculation, a single intraperitoneal EdU injection was administered at E12.5 or E14.5, embryos were fixed after 24h and the percentage of EdU^+^ cells labeled with Ki67 was calculated. For cell-fate analysis, a single intraperitoneal EdU injection was administered at E12.5 or E14.5, and the number of EdU^+^ cells expressing the indicated cellular markers were analyzed at P13. For HES1 expression analysis during different phases of cell cycle, EdU was injected at E14.5, embryos were fixed after 90 min (S+G2), 8h (early G1) or 14h (late G1) and the percentage of EdU^+^ cells labeled with HES1 was calculated. EdU was labeled using Click-iT^®^ EdU imaging kits with alexa 555 or 647 following the manufacturer’s instructions (#C10638 or #C10340, Thermo Fisher Scientific). Briefly, brain sections were post-fixed with 3.7% formaldehyde and permeabilized with 0.5% Triton X-100 in PBS. The Click-iT^®^ reaction cocktail was then added, and the sections incubated in the dark for 30 minutes at room temperature. After the washes, the brain sections were stained for the different proteins of interest using the immunohistochemistry protocol previously described.

### Expression Constructs

pCAG-mRFP (plasmid #32600, Addgene) was deposited by Anna-Katerina Hadjantonakis^57^. pCAG-CRE (plasmid #13775, Addgene) was generated by Connie Cepko^58^. pCS2 NOTCH1 Full Length-6MT (pCS2-*Notch1-FL*, plasmid #41728, Addgene) was deposited by Raphael Kopan^59^. pCAG-Brn-DBD-EnR (plasmid #19714, Addgene) and pCAG-Brn2 (plasmid #19711, Addgene) were generated by Connie Cepko^60^. pCMV6 Delta1 Full Length Myc-DDK was obtained from Origene (pCMV6-*Dll1-FL*, #MR226161). pBlue-hASPM(WT) was generated by Byoung-Il Bae.

### *In utero* electroporation in mice

The protocol used here was adapted from Saito and Nakatsuji^61^. For electroporation of embryonic day 12.5 (E12.5) embryos, five 50 ms pulses of 30 V were delivered (E14.5 – 80ms, 40V). Embryos were left to develop *in utero* for the indicated time or females were allowed to give birth and pups were euthanized at the indicated age.

### *In utero* electroporation in ferrets

The protocol used here followed the established method described by Kawasaki et al.^62^. Briefly, pregnant ferrets were anesthetized with ketamine, and their body temperature was monitored and maintained using a bair hugger warmer. Anesthesia was then continued with isoflurane at 1-2%. Respiratory rate was closely observed, and analgesic solution was injected subcutaneously (Buprenorphine, 0.001 mg/Kg). For E35 electroporation five 50 ms pulses of 100 V were delivered. The embryos were left to develop *in utero* for 48h.

### Luciferase assay

*Ensembl* predicted regulatory region including the POU domain binding site on *Aspm* promoter/enhance (sequence from 5’-GAAAAAGTGGGCAGTAACTCGC-3’ to 5’-CAACCTTTCCCTGAGGACGATC-3’) was synthesized by Twist Bioscience and cloned into pGL3-Basic Luciferase Reporter vector (Promega) using the Gibson assembly method. 150 ng of both pRL-CMV and pGL3-*Aspm* vectors were co-transfected with 0, 100, 200, 400, or 800 ng of either pCAG-Brn2 vector or pCAG-Brn-DBD-EnR. HEK 293 cells were seeded into 96-well plates at 4,000 cells per well and transfected the following day with PEI Prime linear polyethylenimine transfection reagent (Sigma) at a 1:3 ratio of DNA:PEI. Cells were harvested 48 h after transfection and luciferase activity was measured using the Dual-Glo Luciferase Assay system (Promega) on a FlexStation*®* 3 plate reader. Firefly luciferase activity was normalized to *Renilla* luciferase activity.

### Chromatin Immunoprecipitation (ChIP-qPCR)

The interaction of BRN1/2 with the indicated genes was studied by ChIP-qPCR in E14.5 wild-type brains using the ChIP assay kit following the manufacturer’s instructions (Millipore). Immunoprecipitation was performed with 5 µg of anti-BRN1/2 (PA5-30124 Thermo Fisher Scientific) or rabbit IgG as a negative control. This BRN1/2 antibody recognizes both BRN1 and BRN2 protein sequence. Quantitative PCR (qPCR) was performed with 5 ng of ChIP-enriched genomic DNA in a 20 µl reaction using PowerTrack SYBR Green Master Mix (Thermo Scientific) with readout on a QuantStudio 6 Flex System (Thermo Scientific) using standard cycling parameters. ΔΔC_t_ was calculated using *Gapdh* as a housekeeping gene control. Relative expression was calculated using the equation: relative expression =2^−ΔΔCt^. Three technical replicates of three independent biological replicates were run to account for assay variance. Primer sequences used for qPCR are listed in Supplementary Information_9 (SI_9). The expression of *Zic1* was used as a positive control^63^.

### Single cell RNA sequencing

#### Tissue preparation and single cell RNA sequencing

Cortex of E12.5 and E14.5 embryos was dissected and dissociated into single cells using the Papain Dissociation System (#LK0031500, Worthington Biochemical). Individual collected cells were processed for scRNAseq using. Cell counts and viability were determined using the Cell Countess II with Trypan Blue. A maximum volume of 43.3 µl/sample was used for processing to target up to 10,000 cells. Cells were combined with RT reagents and loaded onto 10X Next GEM Chip G along with 3’ v3.1 gel beads. The NextGEM protocol was run on the 10X Chromium Controller to create GEMs, composed of a single cell, gel bead with unique barcode and UMI primer, and RT reagents. 100 µl of emulsion is retrieved from the chip and incubated (45 min at 53°C, 5 min at 85°C, cool to 4°C), generating barcoded cDNA from each cell. The GEMs were broken using Recovery Agent and cDNA cleaned, following manufacturer’s instructions using MyOne SILANE beads. cDNA was amplified for 11 cycles (3 min at 98°C, 11 cycle: 15 sec at 98°C, 20 sec at 63°C, 1 min at 72°C; 1 min at 72°C, cool to 4°C). Samples were cleaned using 0.6X SPRIselect beads. Quality control (QC) was completed using Qubit and Bioanalyzer to determine size and concentrations. 10 µl of amplified cDNA was carried into library prep. Fragmentation, end repair and A-tailing were completed (5 min at 32°C, 30 min at 65°C, cool to 4°C), and samples were cleaned up using double sided size selection (0.6X, 0.8X) with SPRIselect beads. Adaptor ligation (15 min at 20°C, cool to 4°C), 0.8X cleanup and amplification were performed by PCR using unique i7 index sequences. Libraries underwent a final cleanup using double sided size selection (0.6X, 0.8X) with SPRIselect beads. Library QC was performed using Qubit, Bioanalyzer and KAPA library quantification qPCR kit. Libraries were sequenced on the Illumina NovaSeq 6000 using v1.5 kits, targeting 50K reads/cell, at read lengths of 28 (R1), 8 (i7), 91 (R2). Demultiplexing and FASTQ generation was completed using Illumina’s BaseSpace software. Data were aligned to the mouse reference genome (refdata-gex-mm10-2020-A) publicly available at the 10X genomics’ website using the Cell Ranger pipeline. Although we started with seven *Brn1/2-cKO* and seven wild-type littermates of both sexes the genotype for one of the wild-type samples could not be confirmed and therefore this sample was removed from the analysis.

#### Single cell RNA sequencing analysis

Briefly, the 10x data were manually aggregated to create a comprehensive data set and subjected to additional QCing. Raw counts were aggregated together for input into the Monocle3 R/Bioconductor platform and preprocessing performed as previously described^64^. Batch correction was done using the aligned_cds function within Monocle3 for the purpose of visualization^65^. Dimensionality reduction and visualization for the aggregate 10x data were performed using Uniform Manifold Approximation and Projection (UMAP)^66^ for all cells passing QC. Briefly the first 50 principle components of the log10(CPT+1) was used as input for the monocle3 implementation of the UMAP algorithm with the following additional parameters: max_components = 3, metric = ’’cosine’’. Initial cell clusters were called using the louvain algorithm with manual annotation using marker genes. The progenitor and neuronal lineages were subset from the main data using this initial clustering. A new embedding was learned for this subset and cell with ambiguous identity were removed. Although we sequenced seven *Brn1/2-cKO* and seven wild-type littermates of both sexes (total cells = 161,000) the genotype for one of the wild-type samples could not be confirmed and therefore this sample was removed from the subsequent analysis (total cells = 149,712). AP (28% of cells), BP (6.7% of cells), Neurons (31% of cells), DL and UL signatures scores were calculated using the AddModuleScore function from Seurat with cell type signatures from Telley et al.^26,27^ and refined the combination of the results of the clustering and the module Scores with additional marker genes^26–29,67^.TRIcycle analysis was performed as previously described^37^. Final cell types were called using a combination of cluster analysis and cell marker genes^26–29^. All differential expression tests were performed across all expressed genes using the Monocle3 VGAM likelihood ratio test with batch and conditions included in the model formula^64^. Additional statistical tests were carried out in R/Bioconductor. Unpaired t-test was used to compare the total cell types per age and genotype. Wilcoxon rank sum test was used for TRIcycle cell cycle phase analysis and Pearson’s Chi-squared test for given probabilities was used for cell proportion analysis. The heatmaps and dotplots were generated with ggplot2 using the row normalized nonzero means. The signature score heatmaps were generated by averaging the DL signature and UL signature scores within control DLNs, ULNs, and all *Brn1/2-cKO* neurons at each age. To generate the correlation plots, the control ULNs, the control DLNs, and all *Brn1/2-cKO* neurons at each age were pseudo-bulked and the Pearson correlation between each group was calculated within each age. The DE genes were selected using the following thresholds: standard error < 0.15, q value ≤ 0.001 for neurons, AP and BP, and q value ≤ 0.05 for all progenitors (AP and BP together). The ribosomal and mitochondrial genes were removed. GO analysis was performed by the Enrichr API from itapy using the filtered DE lists. The terms of interest were selected for visualization from the significant terms with adjusted q value less than 0.05. The embedding of cynomolgus monkey data was generated by scanpy. Cells with less than 5500 UMIs were excluded. The UMAP was generated with the first 15 principle components and the min_dist of 0.3. AP, BP, UL, and DL signatures scores were calculated using the “score_genes” function. The TRIcycle and DE gene analysis used the same setting as described above. The DE genes were selected by the following criteria: standard error ≤ 0.15, q value ≤ 0.001 for neurons and q value ≤ 0.05 for all progenitors (AP and BP together).

#### Transcriptional waves along pseudotime projection

Only AP cells were selected to learn the pseudotime projection for cell maturity. First, we performed the dimensionality reduction on the control cells. Since one of the sequencing batches (Batch 1) failed to show temporal difference in the latent space, we removed this batch from the following pseudotime analysis. In the latent space of the remaining control data, the second and third principal components best describe the temporal difference and explain a significant amount of variance. Therefore, we fitted a principal curve based on these two PCs using prinPY and projected the cells back into the fitted trajectory to define the maturity pseudotime. The side with highest expression of *Sox2* and *Hmga2* was selected as the beginning as previously described^27^ and the pseudotime score was normalized between 0 to 1. For the *Brn1/2-cKO* cells, we used KDtree to search for their nearest neighbor among control cells in the latent space and assigned the *Brn1/2-cKO* cells with the same pseudotime score as its nearest control neighbor. We then visualized the previously described temporal wave genes using Seaborn heatmap. The cells were aligned according to their pseudotime score and the heatmap was smoothed by the UnivariateSpline. We have confirmed that the control and *Brn1/2-cKO* cells from Batch 1 showed similar average age-related expression pattern compared with the other batches within the same genotype for the vast majority of the genes used to define the transcriptional waves. The Fisher’s Exact test was used to measure the enrichment of the selected GO terms in each wave and the heatmap was colored by the -log10(p-value) from the statistical test.

### Imaging

All immunohistochemistry and ISH RNAscope data were acquired in an equivalent latero-medial level of the cortex from at least 3 independent animals per condition or genotype. Images were captured using a Zeiss LSM 800 confocal microscope. Maximum intensity projections were generated in ImageJ or Imaris software. In all fluorescence microscopy figures, different channels of image series were represented in pseudocolor, and contrast and brightness were adjusted manually using ImageJ or Adobe Photoshop software. All markers were counterstained with DAPI to allow visualization of overall cellular density.

### Histological analysis

A series of *z*-stack (depth of 10 μm) confocal tiled images were used for cell quantification; 150 μm-wide images (embryonic brains) or 300 μm-wide images (postnatal brains) comprising the entire extension of the cortex were analyzed. All quantification were made in an equivalent latero-medial level of the cortex from at least 3 independent animals per condition or genotype. For cortical thickness quantifications, P13 postnatal brains were sectioned coronally at 60 µm and sections sequentially collected over 10 wells. Each well had a full representation of the brain. Three equivalent midbrain sections were imaged and quantified per mice per genotype. The midbrain sections were classified in reference to their proximity to the ventricle, hippocampus, and corpus callosum/midline (1^st^ section ∼Interaural 3.46mm; Bregma -0.34mm; 2^nd^ section ∼Interaural 2.58mm; Bregma -1.22mm; 3^rd^ section ∼Interaural 1.74mm; Bregma -2.06mm).The number of cells expressing the markers of interest was quantified in radial sections of the same latero-medial level (determined as mentioned above) and adjusted per area. For the IUE analysis, to divide the cortex into longitudinal bins, a rectangle was drawn from the border of the white matter with layer VI to the pial surface. This rectangle was then divided into ten equal bins and the number of cells within each bin was quantified. To quantify ISH-RNAscope data, all punctate dots or all punctate dots in RFP^+^ cells were counted in 50 μm-wide images comprising the entire extension of the cortex. *Notch1* mRNA levels in the ferret *in utero* electroporated cells were quantified in all RFP^+^ cells in the area comprising 50 µm from the VZ. For the centriole number analysis, a z-stack of 10 sections (0.5μm each) was selected for all analyzed cells and the number of centrioles overlapping or in the very close proximity of the g-TUBULIN staining were counted. Since *Brn1/2-cKO* show a 20% reduction of total cell number and cortical thickness, the quantifications were normalized to the cortical thickness of each analyzed section in controls and mutants.

### Statistical analysis

Statistical analyses of the immunohistochemistry and ISH-RNAscope experiments were done using GraphPad Prism software. Statistically significant differences were assessed by Student’s unpaired t-test or one-way analysis of variance (one-way ANOVA – Dunnett’s multiple comparisons test; F_DFn, DFd_ (DFn: degrees of freedom in the numerator; DFd: degrees of freedom in the denominator)), comparing two or more groups, respectively. To compare the distribution of RFP^+^ cells in the different bins of the cortex for the two different conditions we used two-way analysis of variance (two-way ANOVA – Šídák’s multiple comparisons test; F_DFn, DFd)_. P<0.05 was considered a significant difference. All values represent individual animals mean ± standard error of the mean (SEM). The statistical test used, and the statistical significance are indicated in figure legends. Statistical analyses of the scRNAseq experiments were done using Monocle 3 and R/Bioconductor. Monocle3 VGAM likelihood ratio test was used for the differential expressed gene analysis. Unpaired t-test was used to compare the total cell types per age and genotype. Wilcoxon rank sum test was used for TRIcycle cell cycle phase analysis and Pearson’s Chi-squared test for given probabilities was used for cell proportion analysis.

## Supporting information

Supplementary Information_revised

## Acknowledgments

We thank members of the Müller and Kolodkin laboratories for suggestions, reagents and technical help; Alex Kolodkin, Jakub Ziak and Cristina Gil-Sanz for critical comments on the manuscript; Caiying Guo for help in generating the *Brn1* floxed mice; Amanda Maxwell, Caroline Garrett and Eric Hutchinson for help with ferrets; Michele Pucak for imaging assistance; Linda Orzolek and Tyler J. Creamer for help with scRNAseq; Anna-Katerina Hadjantonakis, Connie Cepko and Raphael Kopan for plasmids; Ryoichiro Kageyama for the HES1 antibody; Andrew Holland for the γ-TUBULIN and Centrin antibodies. We are grateful to Michelle Monroe, Tajma Smith, Kaiping Zhang, Femi Cleola Villamor and Trinity Walker for assistance with mouse maintenance and genotyping.

## Author contributions

Conceptualization: SB, UM; Methodology: SB, GSO’B, YX, LG, KN, RV, UM; Investigation: SB, GSO’B, YX, UM; Validation: SB, GSO’B, YX, UM; Visualization: SB; Formal analysis: SB, GSO’B, YX, UM; Data curation: SB, GSO’B, YX; Resources: KN, B.-I.B, RSS, CAW, UM; Supervision: UM; Project administration: UM; Funding acquisition: UM; Writing - original draft: SB, UM; Writing - review & editing: SB, YX, LG, KN, RV, B.-I.B, RSS, CAW, GSO’B, UM.

## Data and materials availability

All data are available in the main text or the supplementary information. Correspondence and requests for materials should be addressed to UM.

## Extended Data

**Extended Data Fig. 1.**
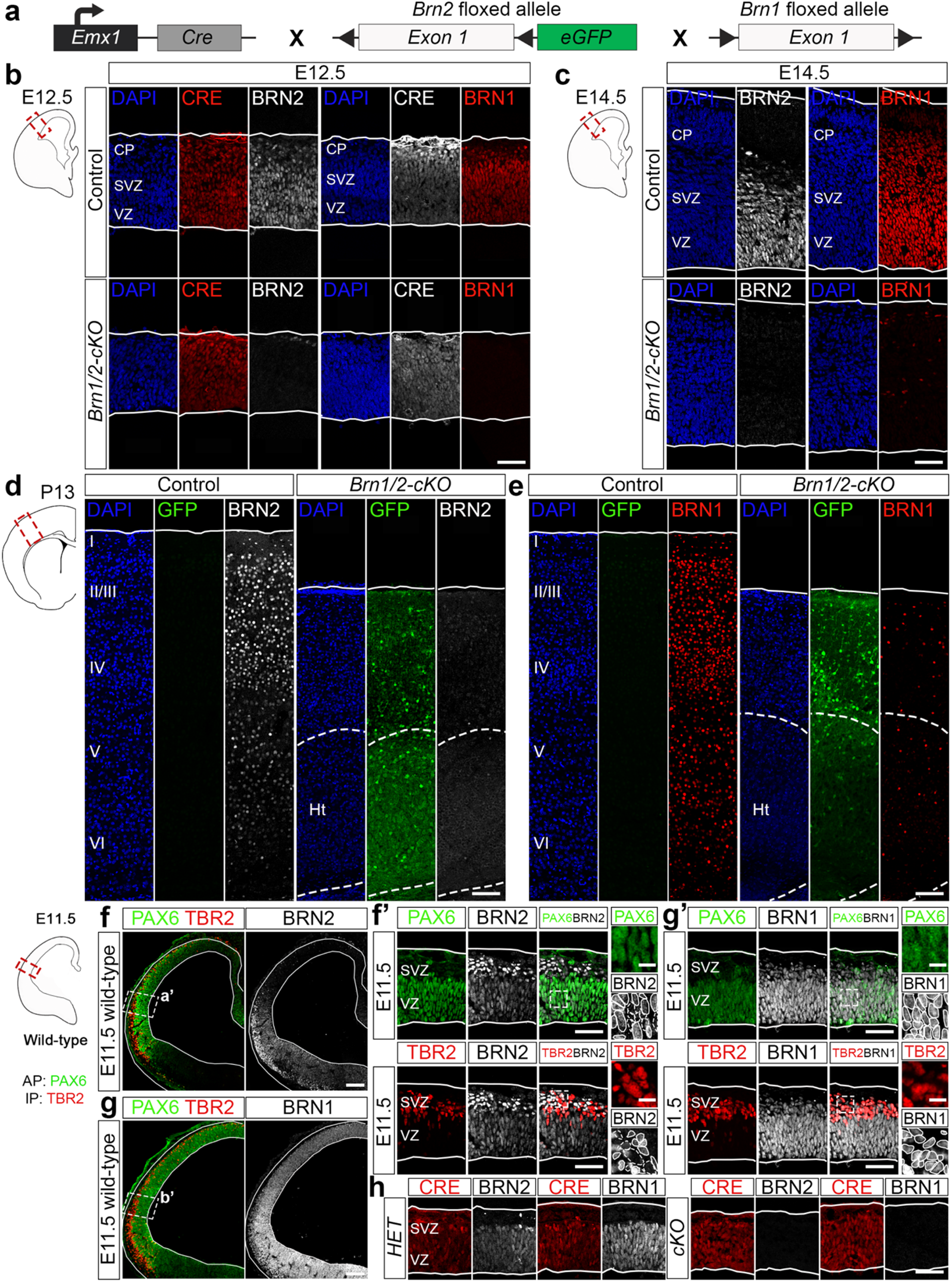
BRN1 and BRN2 expression. **a**, Diagram of experimental strategy. *Emx1-Cre* mice were crossed with *Brn1^fl/fl^*;*Brn2^fl/fl^* mice to inactivate *Brn1* and *Brn2* in the dorsal telencephalon. **b, c,** DAPI staining (blue) and BRN2 (grey), CRE (E12.5, red), BRN1 (red) and CRE (E12.5, grey) immunolabeling in cortical sections of control and *Brn1/2-cKO* mice at E12.5 (b) and E14.5 (c). Low and top lines represent the limits of the VZ and CP, respectively. Scale bar: 50 µm. **d, e,** DAPI staining (blue) and GFP (green), BRN2 (d, grey) and BRN1 (e, red) immunolabeling in cortical sections of control and *Brn1/2-cKO* mice at P13. BRN1 is also expressed in interneurons and therefore those cells, that are not recombined in *Emx1-Cre* mice, remain positive for BRN1 in the cortex of *Brn1/2-cKO* mice at P13 (e). Neocortical cell layers I-VI are indicated. Lines represent the limits of the cortical plate; Dashed lines outline the cortical heterotopia (Ht). Scale bar: 100 µm. **f, g,** Overview of E11.5 wild-type brains labeled for DAPI (blue), PAX6 (green), TBR2 (red), BRN2 (f, grey) and BRN1 (g, grey). Meningeal and ventricular border indicated by white lines. Scale bar: 100 µm. **f’, g’,** Higher magnification view for TBR2 (red), PAX6 (green), BRN2 (f’, grey) or BRN1 (g’, grey) immunolabeling in the squared area highlighted in f or g, respectively. Low and top lines represent the limits of the VZ and CP, respectively. Boxed area shown at higher magnification on the rightmost panels. TBR2^+^ and PAX6^+^ cells expressing or lacking BRN1/2 expression are outlined by lines or dashed lines, respectively. Scale bars: 50 µm (lower magnification), 10 µm (higher magnification). **h,** CRE (red), BRN2 and BRN1 (grey) immunolabeling in cortical sections of control heterozygous and *Brn1/2-cKO* mice at E11.5. Low and top lines represent the limits of the VZ and CP, respectively.

**Extended Data Fig. 2.**
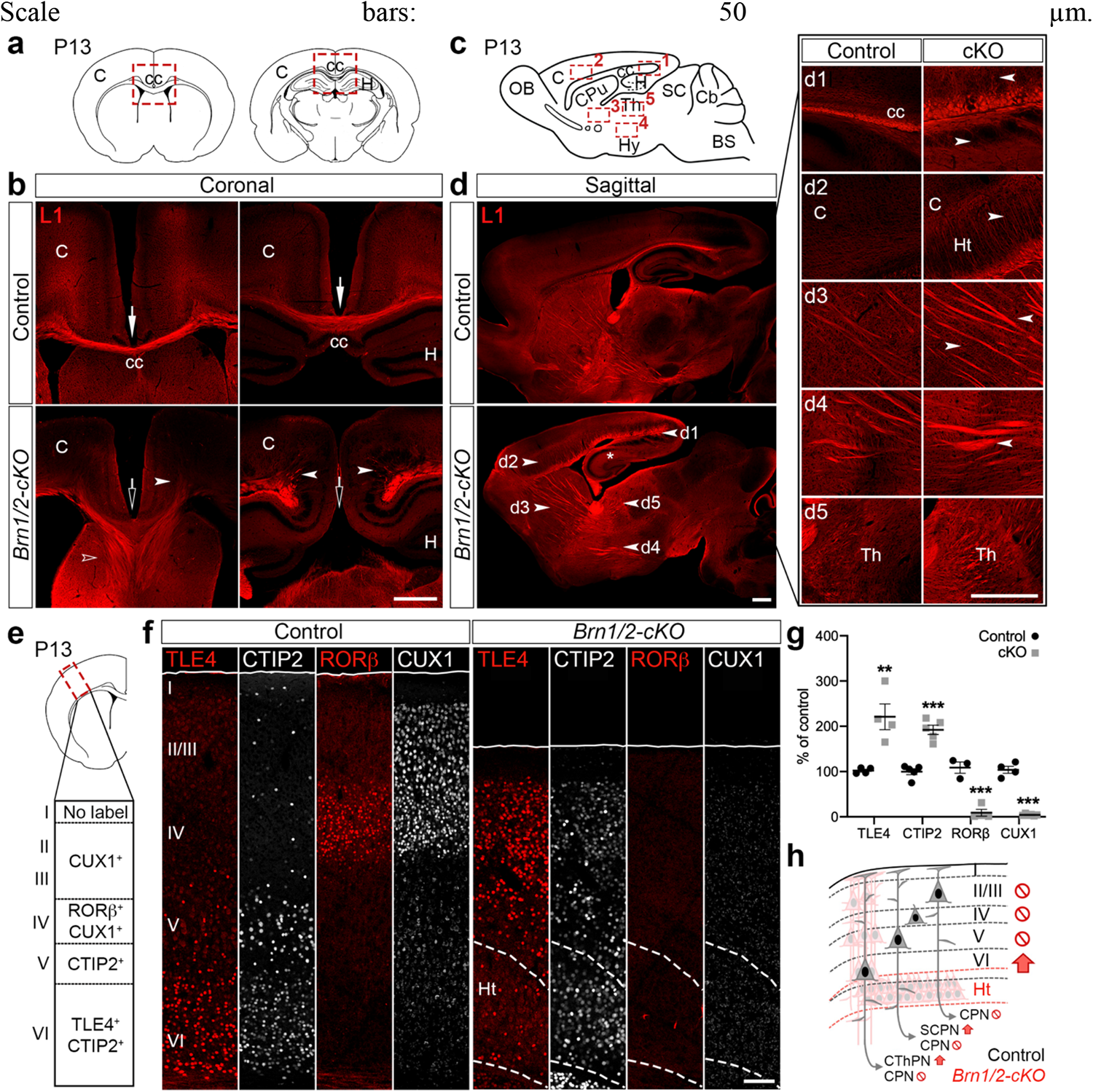
BRN1/2 are essential for proper neuronal specification and circuit development in the neocortex. **a**, Schematic of coronal brain sections at two histological levels highlighting the corpus callosum (cc). C, cortex; H, hippocampus. **b,** Cc of control and *Brn1/2-cKO* mice at different histological levels (500µm intervals) analyzed by L1 immunolabeling (red) at P13. Open arrows indicate absent cc in *Brn1/2-cKO* mice. Open arrowhead shows the abnormal ventral misrouting of L1^+^ projections in *Brn1/2-cKO* mice. Arrowheads indicate misrouting and abnormal defasciculation of L1^+^ neurons in *Brn1/2-cKO* mice. **c,** Schematic of a midsagittal brain section highlighting the cc (1), cortex (C, 2), subcortical projections (3), hypothalamus (Hy, 4) and thalamus (Th, 5). OB, olfactory bulb; SC, superior colliculus; Cb, cerebellum; BS, brain stem. **d,** Neuronal projections of control and *Brn1/2-cKO* mice in midsagittal brain sections analyzed by L1 immunolabeling at P13. Arrowheads point to morphological changes in the *Brn1/2-cKO* brains. Numbers indicate areas shown in panel d1-d5. d1, absent cc; d2, cortical neuronal heterotopia (Ht); d3, more L1^+^ neuronal fibers projecting to subcortical areas; d4, more disorganized L1^+^ neuronal fibers projecting to subcortical areas; d5, more L1^+^ neuronal fibers targeting the thalamus; *disorganized hippocampus as previously described^18^. **e,** Schematic of a coronal brain section highlighting the somatosensory cortex analyzed at P13 by immunolabeling of the indicated molecular markers. **f, g,** Cortical layers of control and *Brn1/2-cKO* mice analyzed by TLE4 (layer VI, red), CTIP2 (layer VI, V and interneurons, grey), RORβ (layer IV, red) and CUX1 (layer IV and II/III, grey) immunolabeling (TLE4: n=4 mice per group, CTIP2: n=5 mice per group, RORβ: n=3 CT, n=4 *cKO* mice, CUX1: n=4 CT, n=6 *cKO* mice; values are mean ± SEM; unpaired t-test; \*\**p<*0.01, ***p<0.001). Neocortical cell layers I-VI are indicated. Lines represent the limits of the cortical plate (CP); dashed lines represent the limits of the cortical heterotopia (Ht). **h,** Schematic of cortical projection neurons in *Brn1/2-cKO* mice (red) and controls (black). Scale bars: 500 µm (b, d, d1-d5), 100 µm (f).

**Extended Data Fig. 3.**
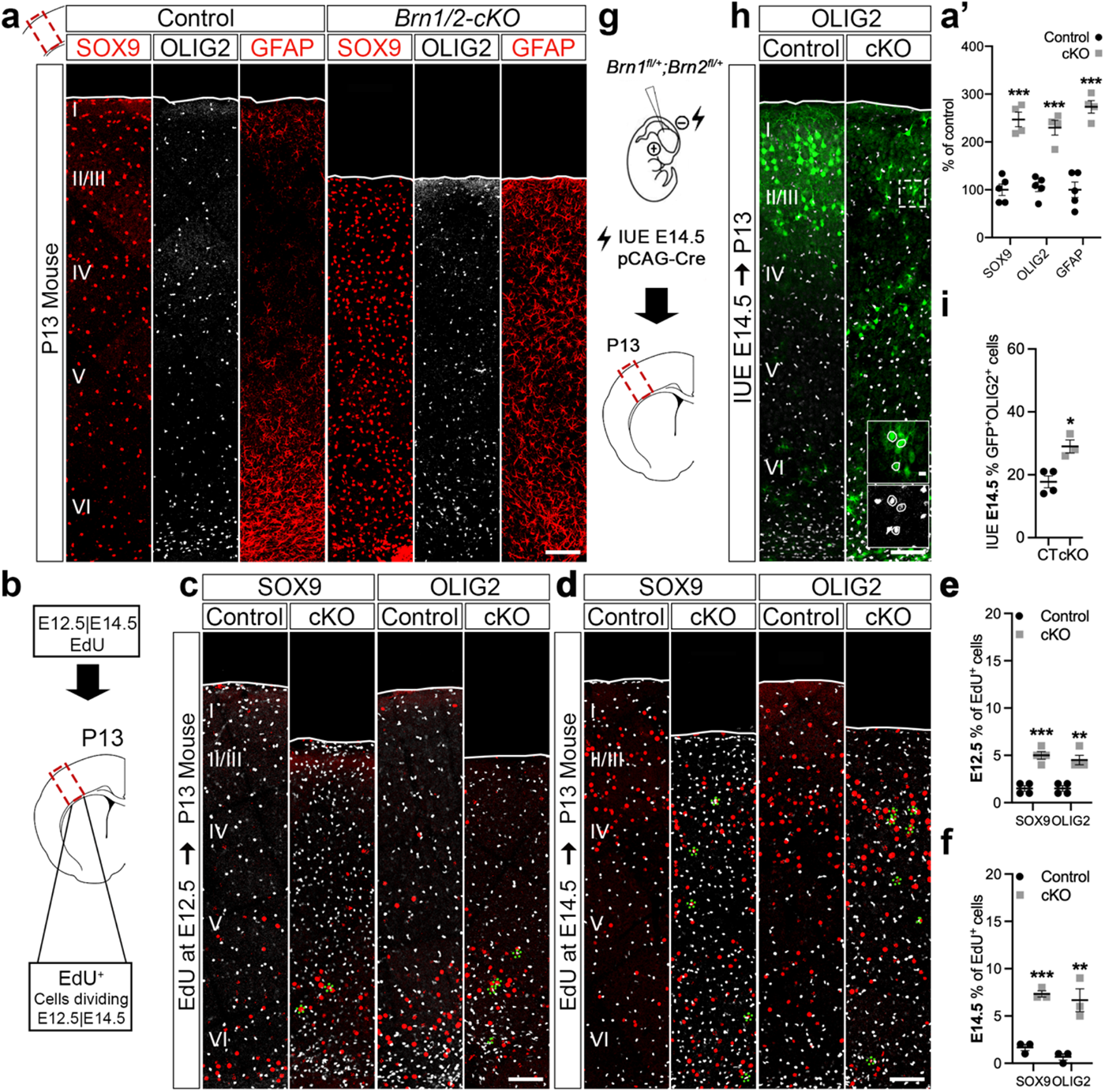
Glial cells are increased in *Brn1/2-cKO*. **a**, a’, SOX9 (red), OLIG2 (grey) and GFAP (red) immunolabeling in cortical sections of control and *Brn1/2-cKO* mice at P13. Neocortical cell layers I-VI are indicated. Lines represent the limits of the cortical plate (n=5 CT, n= 4 *cKO* mice; unpaired t-test). **b,** P13 brains were analyzed by EdU and SOX9 or OLIG2 immunolabeling after intraperitoneal injection of EdU at E12.5 (c) and E14.5 (d). **c-f,** EdU (red) and SOX9 or OLIG2 (grey) immunolabeling in P13 control and *Brn1/2-cKO* cortical sections after EdU injection at E12.5 (c, e) and E14.5 (d, f) (E12.5: n=4 mice per group; E14.5: n=3 mice per group; unpaired t*-*test). Green dotted lines circulating the cells outline cells co-expressing EdU and the indicated glia marker. **g,** IUE in mice from *Brn1^fl/+^*;*Brn2^fl/+^*crossings at E14.5 with the indicated plasmid. **h,** Cell identities of control and *Brn1/2-cKO* condition analyzed by co-immunolabeling for GFP (green; to identify electroporated cells) with OLIG2 (grey) at P13. Lines represent the limits of the CP. Boxed area: higher magnification in inserts. Lines outline cells expressing OLIG2. **i,** GFP^+^ cells expressing OLIG2 at P13 in the control and *Brn1/2-cKO* condition (n=4 CT, n=3 *cKO* mice; unpaired t-test). Values are mean ± SEM; \**p<*0.05, \*\**p<*0.01, ***p<0.001. Top lines represent the limits of the CP. Scale bars: 100 µm (lower magnification), 10 µm (higher magnification).

**Extended Data Fig. 4.**
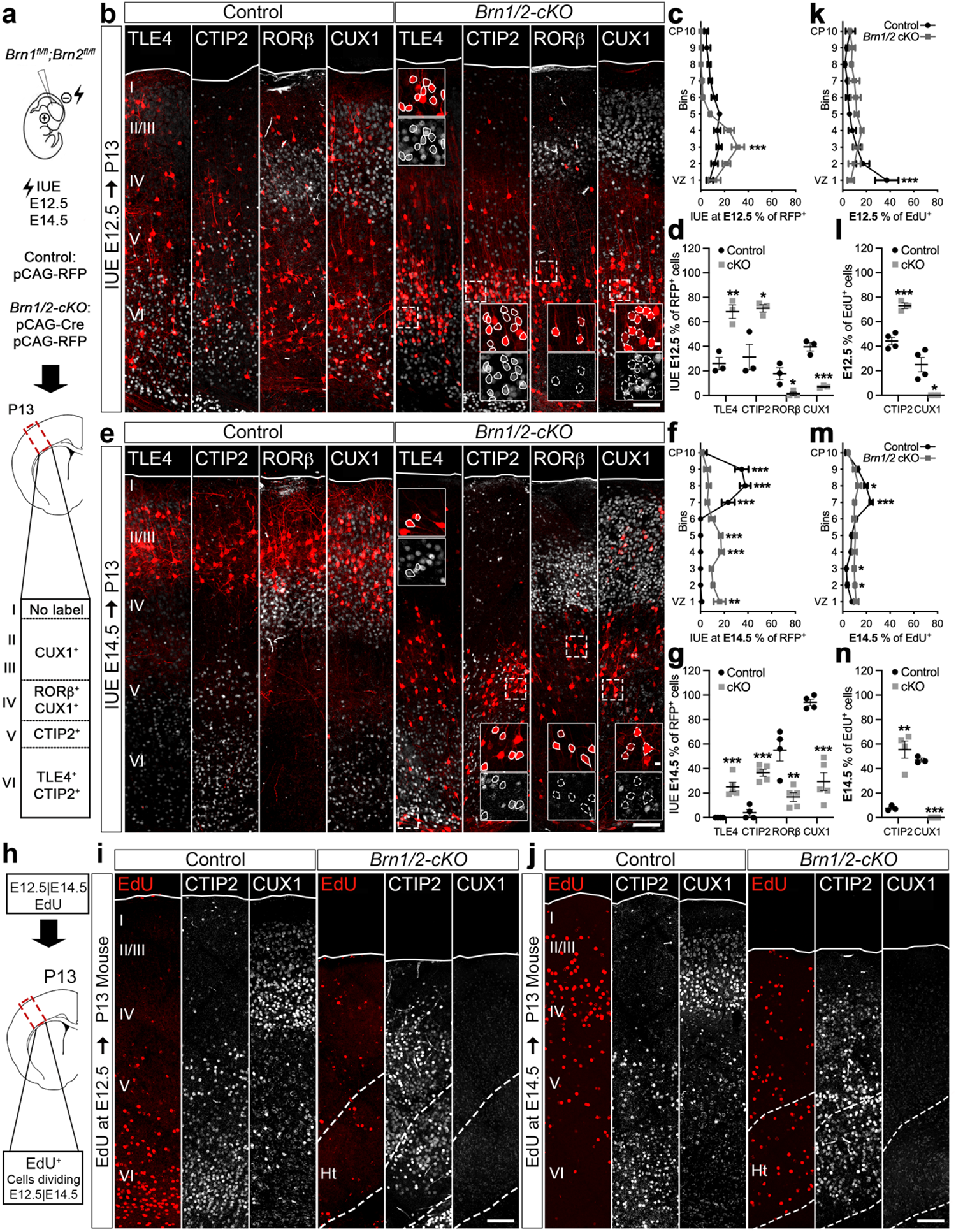
BRN1/2 regulate the competence of progenitor cells to generate ULNs. **a**, IUE in *Brn1^fl/fl^*;*Brn2^fl/fl^* mice at E12.5 and E14.5 with the indicated plasmids. **b, e,** Neuronal identities of control and *Brn1/2-cKO* condition analyzed by co-immunolabeling of RFP (red) to identify electroporated cells with TLE4, CTIP2, RORβ and CUX1 (grey) at P13 after IUE at E12.5 (b) or E14.5 (e). Lines represent the limits of the CP. Boxed area: higher magnification in inserts. Lines outline cells expressing DLN markers; dashed lines outline cells not expressing ULN markers. **c, f,** Distribution of RFP^+^ neurons in the somatosensory cortex at P13 after IUE at E12.5 (c) and E14.5 (f) (E12.5: n=3 mice per group; E14.5: n=4 CT, n=5 *cKO* mice; two-way ANOVA – Šídák’s multiple comparisons test; E12.5: F_9,60_=5.980; E14.5: F_9,70_=24.90). **d, g,** RFP^+^ neurons expressing the indicated marker genes at P13 after IUE at E12.5 (d) and E14.5 (g) (E12.5: n=3 mice per group, E14.5: n=4 CT, n=5 *cKO* mice; unpaired t*-*test). **h,** P13 brains were analyzed by EdU and CTIP2 or CUX1 immunolabeling after intraperitoneal injection of EdU at E12.5 (i) and E14.5 (j). **k, m,** Distribution of EdU^+^ cells in the somatosensory cortex at P13 after EdU injection at E12.5 (k) and E14.5 (m) (E12.5: n=4 CT, n=3 *cKO* mice; E14.5: n=3 CT, n=4 *cKO* mice; two-way ANOVA – Šídák’s multiple comparisons test; E12.5: F_9,50_=4.323; E14.5: F_9,60_=11.15). **l, n,** EdU^+^ cells expressing the indicated marker genes at P13 after EdU injection at E12.5 (l) and E14.5 (n) (E12.5: n=4 CT, n=3 *cKO* mice; E14.5: n=3 CT, n=4 *cKO* mice; unpaired t*-*test). Cortical neuronal heterotopia (Ht). Values are mean ± SEM; \**p<*0.05, \*\**p<*0.01, ***p<0.001. Top lines represent the limits of the CP. Scale bars: 100 µm (lower magnification), 10 µm (higher magnification).

**Extended Data Fig. 5.**
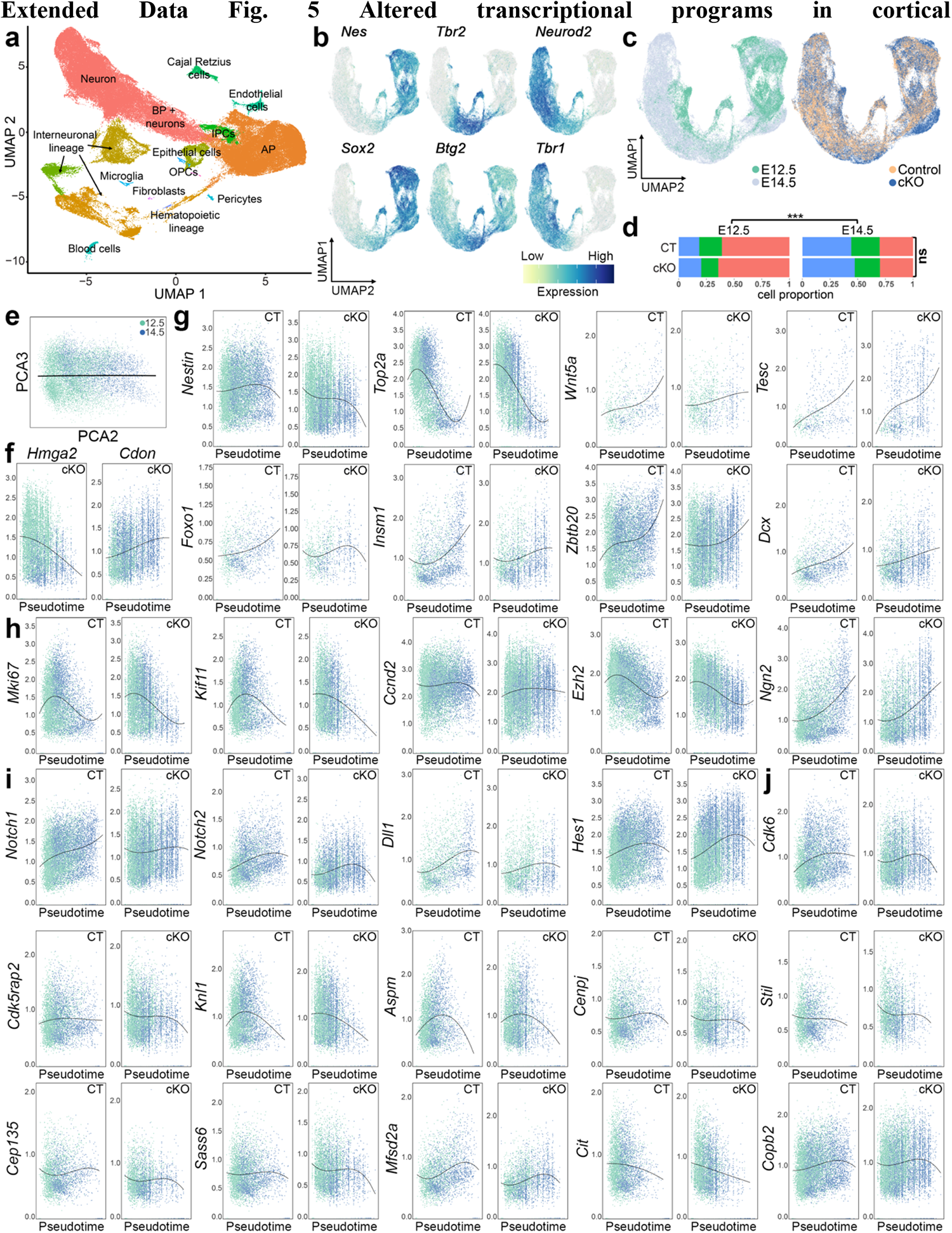
Altered transcriptional programs in cortical progenitors of Brn1/2-cKO mice. **a,** UMAP of scRNAseq data of all cortical cells sequenced at E12.5 and E14.5. BP, basal progenitors; IPCs, intermediate progenitor cells; AP, apical progenitors; OPCs, oligodendrocyte precursor cells. **b,** UMAP of gene signatures for AP (e.g., *Nes*, *Sox2*), BP (e.g., *Tbr2*, *Btg2*) and neurons (e.g., *Neurod2*, *Tbr1*). (**c,**) UMAP of scRNAseq data from control and *Brn1/2-cKO* cortices at E12.5 and E14.5, by age and genotype. **d,** Proportion of cell types by age and genotype (orange = APs, green = BPs, blue = neurons; total cell numbers between genotypes compared by unpaired t*-*test – ns, not significant; cell proportions between ages and different genotypes compared by Pearson’s Chi-squared test – ***p<0.01). **e,** PCA of control AP age identity organization along the pseudotime axis. **f,** Expression of *Hmga2* and *Cdon* in *Brn1/2-cKO* APs along the pseudotime axis. **g,** Examples of different gene expression dynamics in control and *Brn1/2-cKO* APs along the pseudotime axis. **h, i, j,** Expression of cell cycle-associated genes (h), NOTCH signaling-associated genes (i) and microcephaly-associated genes (j) in *Brn1/2-cKO* APs along the pseudotime axis.

**Extended Data Fig. 6.**
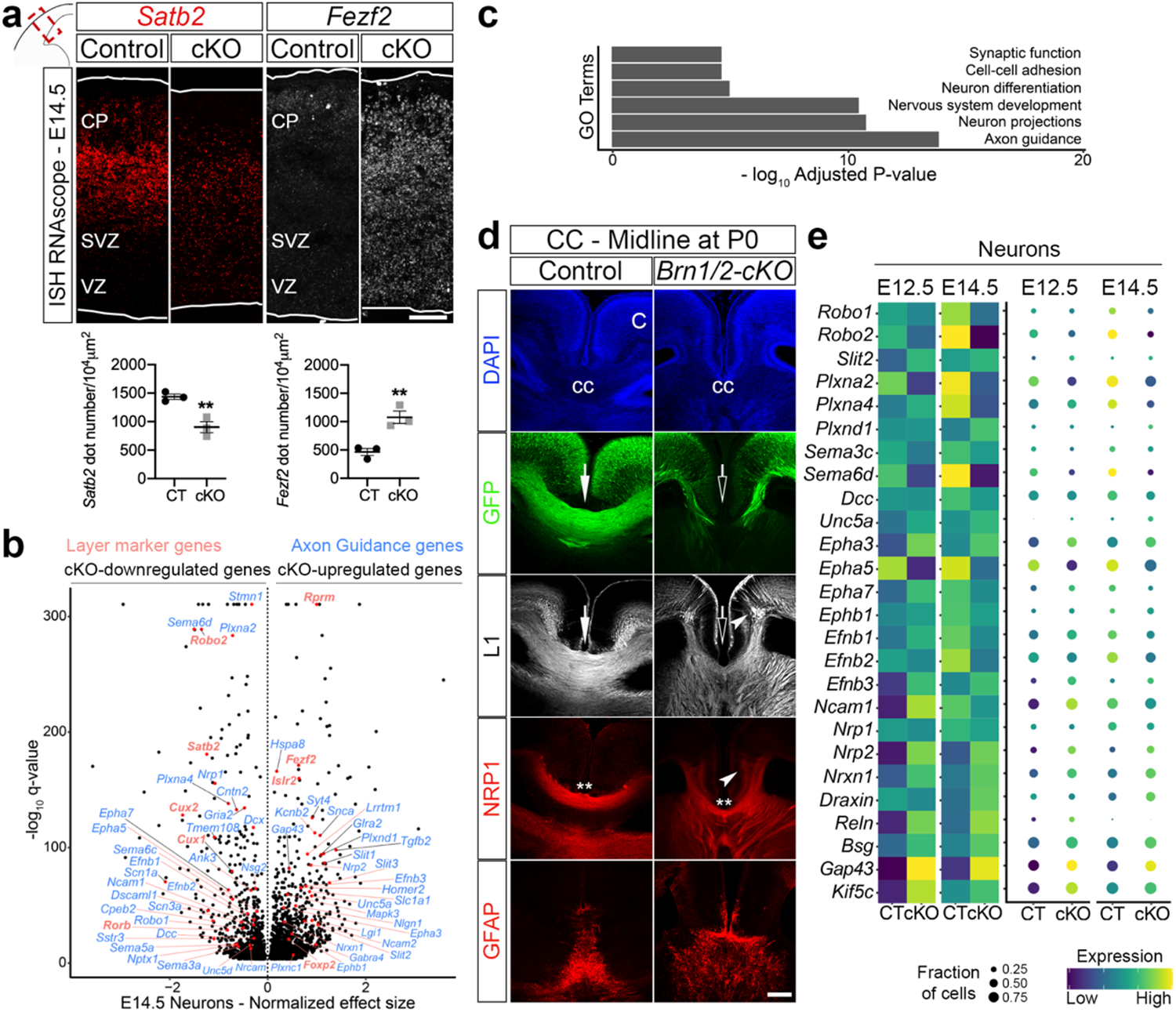
Altered transcriptional programs in *Brn1/2-cKO* cortical neurons. **a**, RNAscope for *Fezf2* and *Satb2* in control and *Brn1/2-cKO* cortical sections at E14.5 (n=3 mice per group; values are mean ± SEM; unpaired t-test; **p<0.01). Low and top lines represent the limits of the VZ and CP, respectively. Scale bar: 50 µm. **b,** Volcano plot: genes differentially expressed between *Brn1/2-cKO* and control neurons at E14.5 highlighted for layer marker and axon guidance genes (Monocle3 VGAM test; SD=0.15; q<0.001; SI_2). **c,** Some of the most relevant gene ontology (GO) terms differentially affected in *Brn1/2-cKO* neurons compared to controls at E14.5. **d,** DAPI (blue), GFP (BRN2^+^ callosal axons, green), L1 (callosal axons; grey), NRP1 (pioneer axons; red) and GFAP (glia guidepost cells; red) immunolabeling in control and *Brn1/2-cKO* brains at P0. Open arrows indicate absent corpus callosum (cc) and ventral misrouting of GFP^+^ and L1^+^ projections in *Brn1/2-cKO* mice. Arrowheads indicate misrouting and abnormal defasciculation of L1^+^ and NRP1^+^ neurons in *Brn1/2-cKO* mice. Double asterisks indicate that some NRP1^+^ pioneer axons still cross the midline in *Brn1/2-cKO* mice. C, cortex. Scale bar: 200 µm. **e,** Expression of axon guidance-associated genes in *Brn1/2-cKO* neurons compared to control at E12.5 and E14.5.

**Extended Data Fig. 7.**
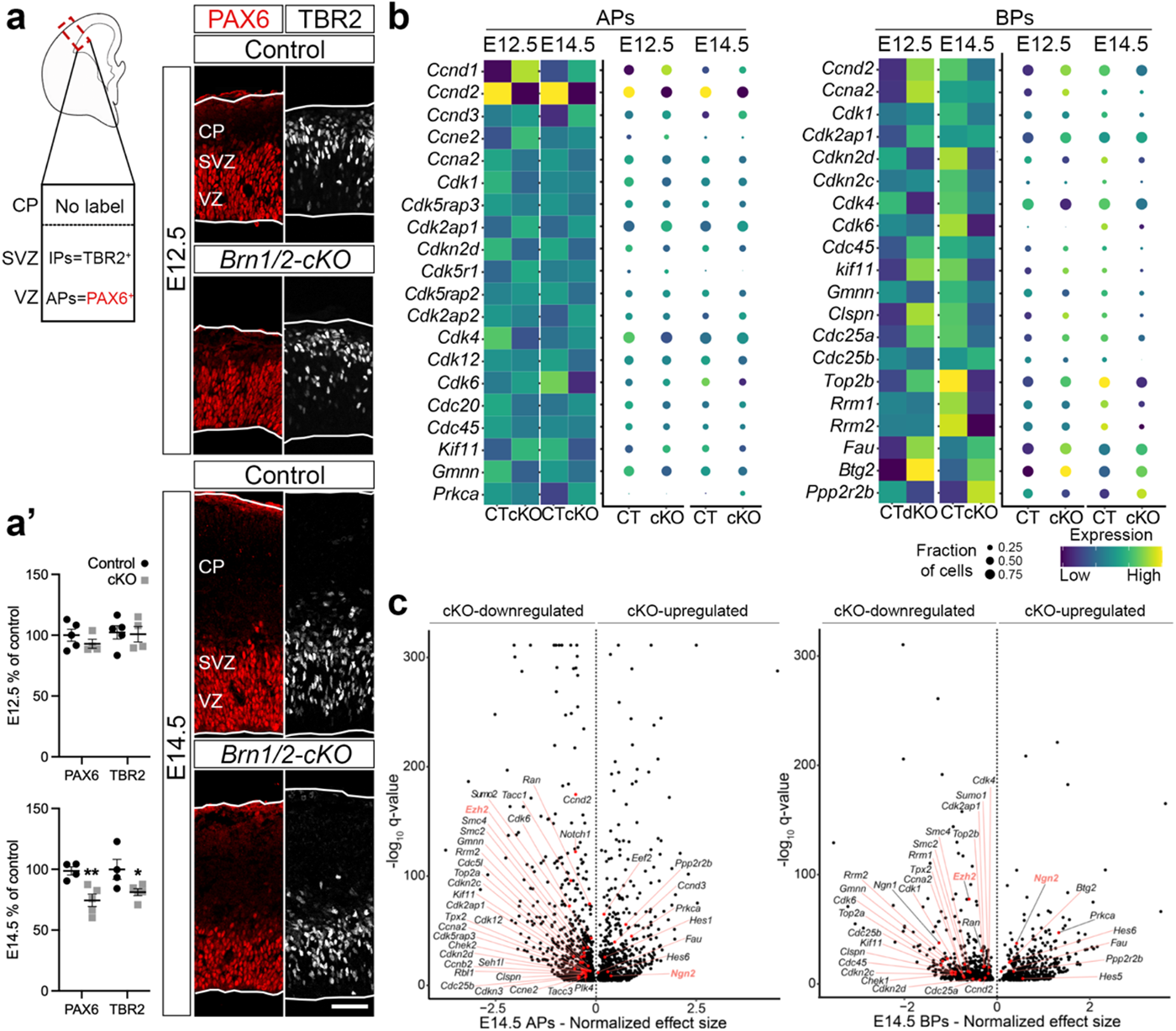
Reduction in proliferation rate and precocious cell cycle exit of cortical progenitors in *Brn1/2-cKO* mice. **a, a’** PAX6 (APs; red) and TBR2 (IPs; grey) immunolabeling in control and *Brn1/2-cKO* cortical sections at E12.5 and E14.5 (E12.5: n=5 CT, n=4 *cKO* mice; E14.5: n=4 CT, n=5 *cKO* mice; values are mean ± SEM; unpaired t*-*test; \**p<*0.05, \*\**p<*0.01). Low and top lines represent the limits of the VZ and CP, respectively. Scale bar: 50 µm. **b,** Expression of cell cycle-associated genes in *Brn1/2-cKO* APs and BPs compared to control at E12.5 and E14.5. **d,** Volcano plots: genes differentially expressed between *Brn1/2-cKO* and control APs and BPs at E14.5 highlighted for cell cycle-associated genes (Monocle3 VGAM test; SD=0.15; q<0.001; SI_3 and SI_4).

**Extended Data Fig. 8.**
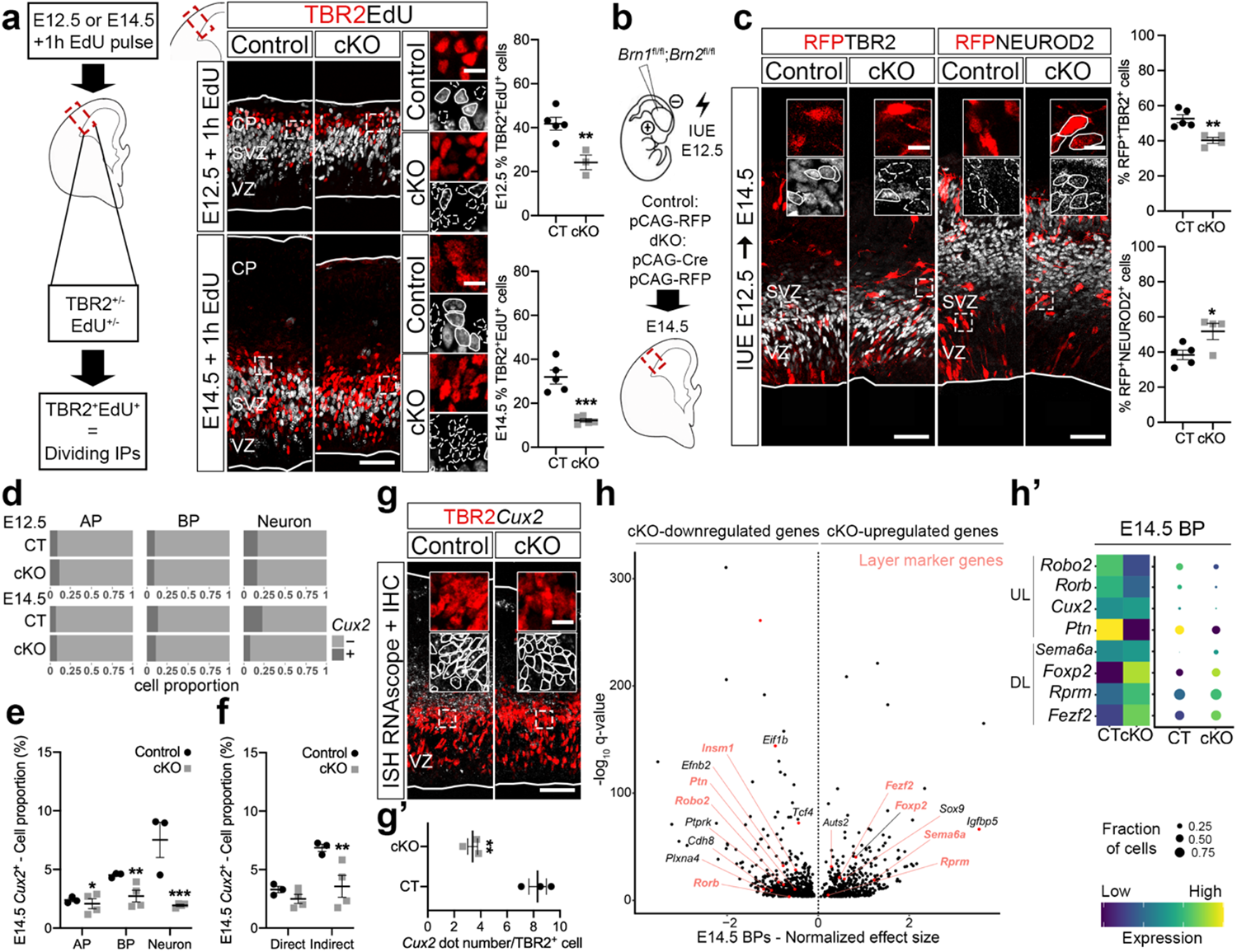
BRN1/2 regulate the balance between direct and indirect neurogenesis affecting the generation of a specific type of BPs. **a**, TBR2 (red) and EdU (grey) immunolabeling in control and *Brn1/2-cKO* cortical sections at E12.5 and E14.5 (E12.5: n=5 CT, n=3 *cKO* mice; E14.5: n=5 mice per group; unpaired t*-*test). Boxed area at higher magnification on the right. Lines and dashed lines circulating cells: expression or absence of EdU, respectively. **b,** IUE in *Brn1^fl/fl^*;*Brn2^fl/fl^* mice at E12.5 with the indicated plasmids. **c,** Cell identities of the control and *Brn1/2-cKO* condition analyzed by co-immunolabeling for RFP (red; to identify electroporated cells) with TBR2 or NEUROD2 (grey) at E14.5 (n=5 CT, n=4 *cKO* mice; unpaired t*-*test). Boxed area: higher magnification in inserts. Lines and dashed lines outlining cells expressing or lacking expression of the indicated marker, respectively. **d,** Proportion of AP, BP and neurons expressing *Cux2* in control and *Brn1/2-cKO* cortices at E12.5 and E14.5. **e,** Proportion of AP, BP and neurons expressing *Cux2* in control and *Brn1/2-cKO* cortices at E14.5 (n ≥ 3 mice per group; Pearson’s Chi-squared test). **f,** Proportion of cells undergoing direct and indirect neurogenesis expressing *Cux2* in control and *Brn1/2-cKO* cortices at E14.5 (n ≥ 3 mice per group; Pearson’s Chi-squared test). **g, g’,** *Cux2* (grey) expression by RNAscope and TBR2 (red) expression by immunolabeling in control and *Brn1/2-cKO* cortical sections at E14.5 (n=3 mice per group; unpaired t*-*test). Boxed area: higher magnification in inserts. Lines oultline TBR2^+^ cells quantified for *Cux2* mRNA expression. **h,** Volcano plot: genes differentially expressed between *Brn1/2-cKO* and control BPs at E14.5 highlighted for layer marker genes (Monocle3 VGAM test; SD=0.15; q<0.001; SI_4). **h’,** Expression of the indicated cortical UL and DL marker genes in control and *Brn1/2-cKO* BPs compared to controls at E14.5. Low and top lines represent the limits of the VZ and CP, respectively. Values are mean ± SEM; \**p<*0.05, \*\**p<*0.01, ***p<0.001. Scale bars: 50 µm (lower magnification), 10 µm (higher magnification).

**Extended Data Fig. 9.**
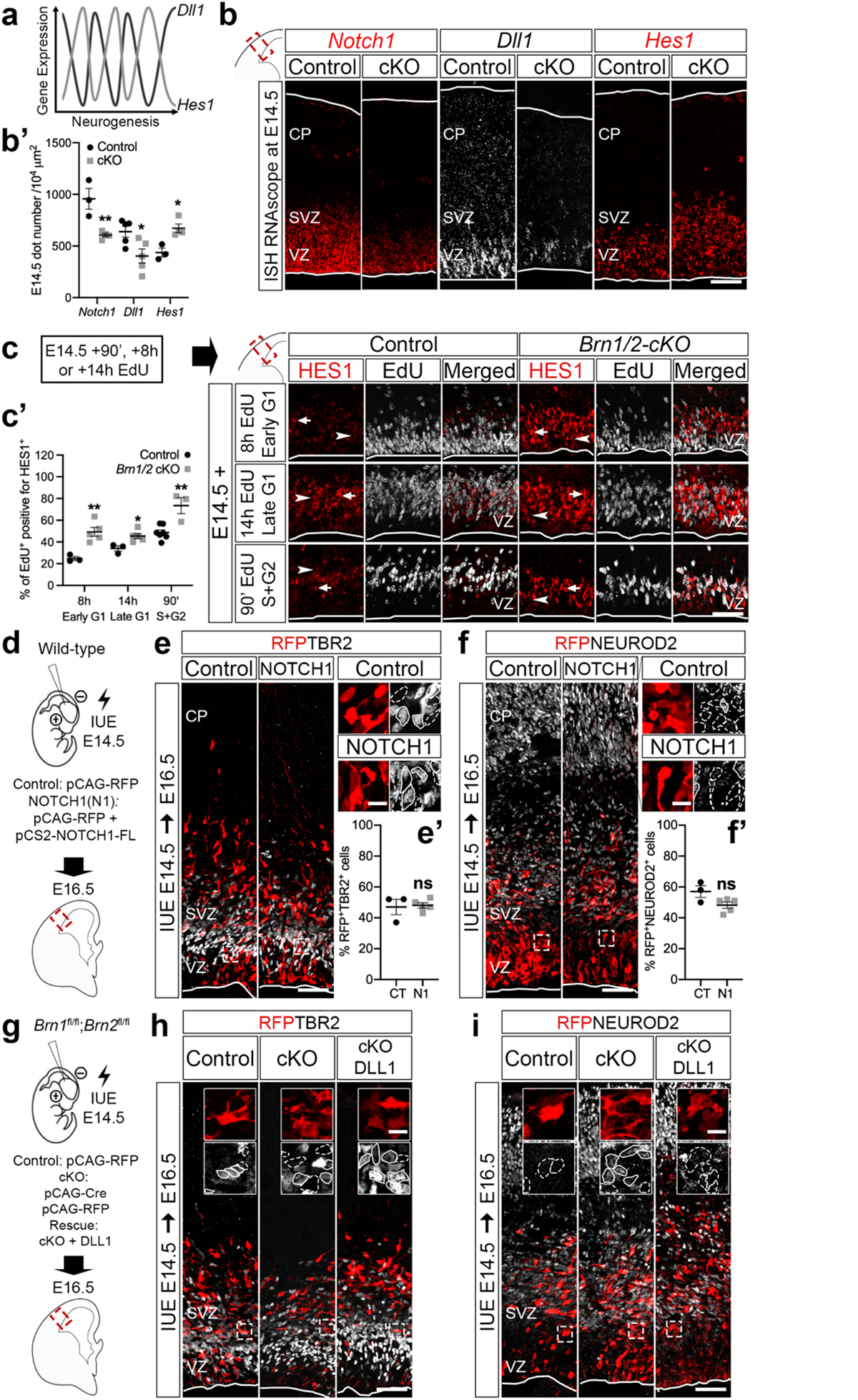
BRN1/2 are essential to maintain NOTCH signaling balanced during neurodevelopment. **a**, Expression of HES1 and DLL1 in neural progenitors is oscillatory, which is critical for the maintenance of neural progenitors^43,44^. **b, b’,** RNAscope for *Notch1* (red), *Dll1* (grey) and *Hes1* (red) in control and *Brn1/2-cKO* cortical sections at E14.5 (*Notch1* and *Hes1*: n=3 CT, n=4 *cKO* mice; *Dll1*: n=5 mice per group; unpaired t*-*test). **c,** E14.5 brains were collected 90’, 8h or 14h after intraperitoneal injection of EdU. HES1 (red) and EdU (grey) immunolabeling in control and *Brn1/2-cKO* cortical sections at early G1 (8h EdU), late G1 (14h EdU) and S+G2 (90’ EdU). Cells expressing HES1 at high and low levels are indicated by arrows and arrowheads, respectively. **c’,** HES1^+^/EdU^+^ cells in control and *Brn1/2-cKO* cortices at early G1 (8h EdU), late G1 (14h EdU) and S+G2 (90’ EdU) (8h and 14h: n=3 CT, n=5 *cKO* mice; 90’: n=7 CT, n=3 *cKO* mice; unpaired t*-*test). **d,** IUE in wild-type mice at E14.5 with the indicated plasmids. **e, f,** Cell identities of the control and NOTCH1 condition analyzed by co-immunolabeling of RFP (red) with TBR2 (e, grey) and NEUROD2 (f, grey) at E16.5. Boxed area at higher magnification on the right. **e’, f’,** RFP^+^ cells expressing TBR2 (e’) and NEUROD2 (f’) (n=3 CT, n=5 N1 mice; unpaired t*-* test). **g,** IUE in *Brn1^fl/fl^*;*Brn2^fl/fl^* mice at E14.5 with the indicated plasmids. **h, i,** Cell identities of the control, *Brn1/2-cKO* and *Brn1/2-cKO*+DLL1 condition analyzed by co-immunolabeling of RFP (red) with TBR2 (h, grey) and NEUROD2 (i, grey) at E16.5. Boxed area: higher magnification in inserts. Lines and dashed lines outlining cells expressing or lacking expression of the indicated marker, respectively. Low and top lines represent the limits of the VZ and CP, respectively. Values are mean ± SEM; ns, not significant; \**p<*0.05, \*\**p<*0.01. Scale bars: 50 µm (lower magnification), 10 µm (higher magnification).

**Extended Data Fig. 10.**
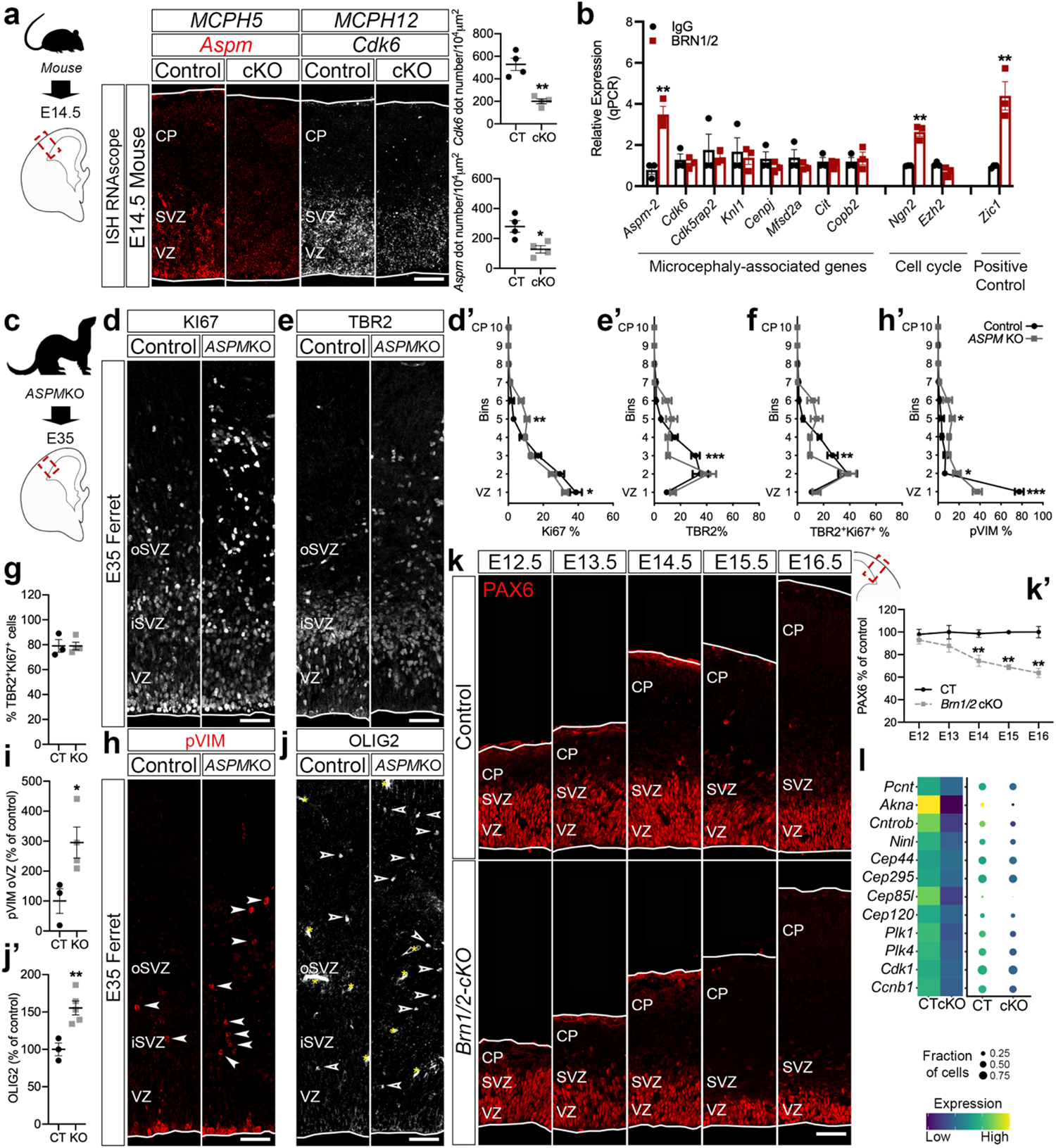
BRN1/2 are required for the expression of microcephaly-associated genes and maintenance of the neuronal progenitor pool. **a**, RNAscope for *Aspm* (red) and *Cdk6* (grey) in control and *Brn1/2-cKO* cortical sections at E14.5 (n=4 mice per group; unpaired t*-*test). **b,** ChIP-qPCR analysis of BRN1/2 binding to the promoters/enhancers of the indicated genes at E14.5 (n=3 per condition; unpaired t-test). **c,** Histological analysis was performed at E35 in cortical sections of control and *ASPM*KO ferrets. **d, e,** Ki67 (d) and TBR2 (e) immunolabeling in cortical sections of control and *ASPM*KO ferrets at E35. **d’, e’, f,** Distribution of Ki67^+^ (d’), TBR2^+^ (e’) and TBR2^+^Ki67^+^ (f) cells in the cortex of control and *ASPM*KO ferrets (Ki67: n=3 CT, n=5 KO ferrets; TBR2 and TBR2^+^Ki67^+^: n=3 CT, n=4 KO ferrets; two-way ANOVA – Šídák’s multiple comparisons test; Ki67: F_9,60_=4.772; TBR2: F_9,50_=5.233; TBR2^+^Ki67^+^: F_9,50_=3.735). **g,** TBR2^+^Ki67^+^ cells in the cortex of control and *ASPM*KO ferrets (n=3 CT, n=4 KO ferrets; unpaired t*-*test). **h,** pVIM immunolabeling in cortical sections of control and *ASPM*KO ferrets at E35. Arrowheads: pVIM^+^ cells outside the VZ. **h’,** Distribution of pVIM^+^ cells in the cortex of control and *ASPM*KO ferrets (n=3 CT, n=4 KO ferrets; two-way ANOVA – Šídák’s multiple comparisons test; F_9,50_=19.02). **i,** pVIM^+^ cells outside of the VZ (oVZ) in control and *ASPM*KO cortices (n=3 CT, n=4 KO ferrets; unpaired t-test). **j, j’,** OLIG2 immunolabeling in cortical sections of control and *ASPM*KO ferrets at E35 (n=3 CT, n=5 KO ferrets; unpaired t-test). Empty arrowheads indicate OLIG2^+^ cells. **k, k’,** PAX6 immunolabeling in control and *Brn1/2-cKO* cortical sections from E12.5 to E16.5 (E12.5: n=6 CT and n=4 *cKO* mice; E13.5: n=5 CT and n=6 *cKO* mice; E14.5: n=4 CT and n=5 *cKO* mice; E15.5: n=5 CT and n=3 *cKO* mice; E16.5: n=5 CT and n=3 *cKO* mice; unpaired t-test). **l,** Expression of centrosome-associated genes in *Brn1/2-cKO* compared to control progenitors at E14.5. Low and top lines represent the limits of the VZ and CP, respectively. Values are mean ± SEM; \**p<*0.05, \*\**p<*0.01, ***p<0.001. Scale bar: 50 µm.

